# Parallelism between phylogeny and ontogeny

**DOI:** 10.1101/2024.06.27.600990

**Authors:** Juraj Bergman, Robert Bakarić, Krunoslav Brčić-Kostić

**Affiliations:** Center for Ecological Dynamics in a Novel Biosphere (ECONOVO), Department of Biology, Aarhus University, Aarhus, Denmark; Section for Ecoinformatics and Biodiversity, Department of Biology, Aarhus University, Aarhus, Denmark; Laboratory for Information and Signal Processing, Department of Electronics, Ruđer Bošković Institute, Zagreb, Croatia; Laboratory for Evolutionary Genetics, Department of Molecular Biology, Ruđer Bošković Institute, Zagreb, Croatia

## Abstract

Haeckel’s biogenetic law, or the recapitulation theory remains a controversial subject to this day. Currently, the modern version of biogenetic law is the hourglass model with its phylotypic period. Importantly, the hourglass model is nothing more than a model of development, and it does not provide any evidence that ontogeny recapitulates phylogeny. However, the hourglass model and biogenetic law are not mutually exclusive, and there are several examples of recapitulation-like processes observable after the phylotypic period of ontogeny. At the level of transcriptomics, all attempts to demonstrate recapitulation failed. Using a novel approach, combining transcriptomics with phylostratigraphy, we demonstrate that recapitulation, or parallelism between phylogeny and ontogeny, exists. We show that the mean indispensability of genes decreases for phylogenetically younger genes, as well as genes expressed during later stages of ontogeny. We also define the ontotypic period of phylogeny, an analog to the phylotypic period of ontogeny. Since it starts from the beginning of phylogeny, it is reasonable to hypothesize that recapitulation starts from the phylotypic period. We conclude that parallelism, or recapitulation, is explainable by the fact that genes that emerged later in phylogeny have tendencies to be expressed during later stages of ontogeny.

## INTRODUCTION

One of the most influential and controversial concepts in biology is the biogenetic law which considers the relationship between phylogeny and ontogeny. Although there were researchers who studied the connection between phylogeny and ontogeny earlier (Meckel 1811; Muller 1864), the most popular view was articulated in Haeckel’s famous statement that "ontogeny is the short and fast recapitulation of phylogeny” (Haeckel 1866; 1868; 1872). Haeckel assumed that transformations during embryonic development of particular species include some adult features of its ancestors. The recapitulation theory was also a strong argument in favor of common ancestry, a concept introduced by Charles Darwin (Darwin 1859). Although Haeckel’s biogenetic law became very popular, it was also widely criticized. The first criticism came from Karl von Baer who disagreed with the view that embryos could resemble adult forms of ancestral species (von Baer 1828). Most criticisms since the emergence of genetics were based on three issues: 1) recapitulation, as a theory based on comparative morphology and embryology, is not explainable by genetics (Gould 1977; Levit *et al*. 2021); 2) Haeckel used an old and discarded Lamarckian view of evolution (Gould 1977); and 3) Haeckel assumed that embryos showed adult ancestral features during development (Gould 1977; Richardson and Keuck 2002).

These criticisms are not particularly serious. Haeckel was aware that the biogenetic law is a thesis based on comparative morphology and embryology, rather than a law of nature based on quantitative regularities (Haeckel 1866). Although Haeckel in his drawings used a linear representation of evolution, as a devoted Darwinist, he knew that evolution is a branching process. Similarly, the evolutionary series of horses (*Hyracotherium* – *Mesohippus* – *Merychippus* – *Pliohippus - Equus*) are often presented as a linear series, although they evolved by a branching process. It is important to emphasize that during branching evolution there are periods of common evolution for several taxa which implies that recapitulation (but not repetition) is theoretically possible. Therefore, it could be that partial similarity between embryos and ancestral adult features exists. In agreement with this, there are several examples of recapitulation-like processes (Swan 1990; Cohn and Tickle 1999; Lovejoy 2000; Metscher and Ahlberg 2001; Bejder and Hall 2002; Lovejoy *et al*. 2004; Thewissen *et al*. 2006; Nagashima *et al*. 2009; Cooper *et al*. 2014; Botelho *et al*. 2015; Diogo *et al*. 2015; Botelho *et al*. 2017; Thewissen 2018).

It has often been proposed that the modern version of Haeckel’s biogenetic law is the hourglass model with its phylotypic period (Olsson *et al*. 2017). However, this is not correct since the hourglass model is just a model of development, whereas Haeckel’s law emphasizes the connection between phylogeny and ontogeny. There are two contemporary models of development: the funnel (early conservation) (Riedl 1978; Rasmussen 1987), and the hourglass model (Duboule 1994; Raff 1996). According to the hourglass model, early and late periods of development are variable, whereas the middle (phylotypic) period is conserved. The phylotypic period is a period of development when embryos of all species within a phylum are morphologically most similar. This view is also supported by genomic and transcriptomic data. It was shown that the most similar gene expression pattern among different species occurs during the phylotypic period (Hazkani Covo *et al*. 2005; Irie and Sehara-Fujisawa 2007; Kalinka *et al*. 2010; Irie and Kuratani 2011). In addition, it was demonstrated using the transcriptome age index (*TAI*), that the oldest genes are expressed during the phylotypic period of zebrafish (*Danio rerio*) and *Drosophila* development (Domazet-Lošo and Tautz 2010). However, *TAI* is not an adequate parameter to estimate the average age of genes expressed in the transcriptome if the expression and the age of genes are correlated (Supplement 1; Piasecka et al. 2013). It is important to note that the phylotypic period and the models of embryogenesis (hourglass and early-conservation models) do not demonstrate the existence of parallelism between phylogeny and ontogeny, but could be compatible with it (Uesaka *et al*. 2019; Uesaka *et al*. 2021). In order to test whether parallelism between phylogeny and ontogeny exists, we use a novel approach combining genomics, transcriptomics and phylostratigraphy.

## RESULTS

### Parallelism

First, we wanted to detect whether the phenomenon of parallelism between phylogeny and ontogeny exists. Therefore, we compared the dynamics of mean indispensability of genes for the periods of their emergence in phylogeny with the dynamics of mean indispensability of genes for the periods of their expression in ontogeny. To reconstruct the emergence of genes during phylogeny we used the phylostratigraphic approach (Domazet-Lošo *et al*. 2007) which was developed as a working software (Arendsee *et al*. 2019) (Fig. 1). To monitor the expression of genes during ontogeny we used the available transcriptomic data sets (Graveley *et al*. 2011; Li *et al*. 2014; Domazet-Lošo and Tautz 2010; Hu *et al*. 2017). We consider gene indispensability to be proportional to the decrease of organismal fitness due to the loss of function of a gene. Therefore, to provide a quantification of gene indispensability, we consider three parameters related to gene function. We analyzed the housekeeping potential of genes (related to pleiotropy) (Schug *et al*. 2005), the interaction potential of genes (related to epistasis) (Szklarczyk *et al*. 2023) and the *dN/dS* ratio (inversely proportional to selective constraint). We expect genes with higher housekeeping and interaction potential to have higher indispensability as they tend to underlie basic cellular functions and/or affect multiple phenotypes, and thus have higher functional importance. Conversely, we expect genes with higher *dN/dS* ratios to have lower indispensability, as they are likely to have emerged in more recent evolutionary history and are in the early stages of acquiring functionalization, resulting in their lower relevance to organismal fitness.

**Figure 1.**
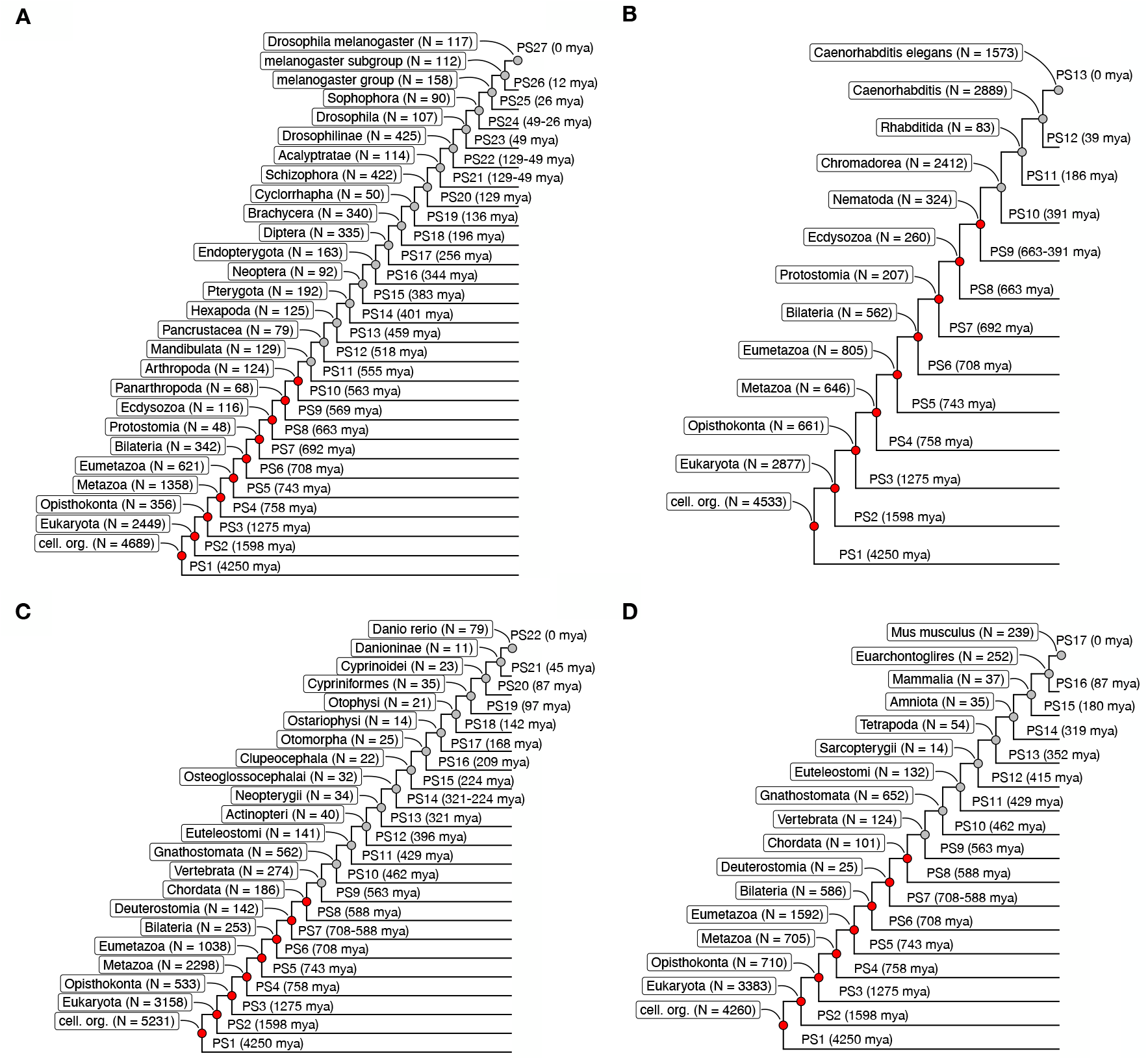
Inferred phylostrata (PS) for analyzed species. A. *Drosophila melanogaster*, B. *Caenorhabditis elegans*, C. *Danio rerio*, and D. *Mus musculus*. The number of genes in each phylostratum (N) is presented in parentheses. Only genes that are expressed during ontogeny are considered. The number associated with each phlyostratum corresponds to its rank, with lower values signifying evolutionarily older phlyostrata and higher values signifying younger phlyostrata. The nodes depicted in red signify the ontotypic period of phylogeny.

Each gene can be classified into a single phylogenetic age category (phylostratum) using the phylostratigraphy approach. Therefore, determining the dynamics of mean gene indispensability during phylogeny is a simple matter of calculating the mean values of indispensability parameters across genes that emerged in each phylostratum. However, before determining dynamics of mean gene indispensability during ontogeny, we first needed to define a subset of functionally expressed genes for each ontogenetic period. This step was necessary as genes are usually expressed across many (and sometimes all) ontogenetic periods, and thus their functional importance across different ontogenetic periods is likely to be obscured if we consider non-zero expression as the only criterion of gene functionality. We therefore consider that a gene is functionally expressed in a specific ontogenetic period if its expression level is higher than a specific cut-off value. This cut-off was defined as a quantile of gene expression determined from the distribution of ontogenetic expression values for each gene. However, it is important to note that this is a very difficult task since it is likely that each gene has its own expression cut-off value which determines its functionality. The problem is even more complicated when one takes into account that most genes are expressed in several periods of ontogeny with possibly different cut-off values for each period. Since these details are not known, we simplify the situation by assuming that in each analysis all genes have the same quantile of expression as a cut-off value. To further address these issues, all analyses were conducted given a wide range of cut-off values, the majority of which resulted in similar indispensability dynamics (Suppl. Figs. 1-16; Suppl. Tables 1-3). In further text, we present an analysis conducted using the 0.1 expression quantile for the cut-off value, as a representative result.

**Table 1.** Ratio of the number of genes in the intersection of genes expressed in the phylotypic stage of ontogeny and the genes that emerged in specific phylogenetic periods, over the corresponding number of genes in the union of the two sets (proportion A). Proportion of the number of genes expressed in the phylotypic stage of ontogeny that emerged in specific phylogenetic periods, out of the total number of genes expressed in the phylotypic stage of ontogeny (proportion B). Proportion of the number of genes expressed in the phylotypic stage of ontogeny that emerged in specific phylogenetic periods, out of the total number of genes that emerged in the corresponding phylogenetic periods (proportion C). Data corresponds to the 0.1 expression quantile for the cut-off value.

|  |  | Blastea | Gastrea | Ontotypic period | BGO |
| --- | --- | --- | --- | --- | --- |
| *Drosophila melanogaster* | A | 54.68%<br>(6,976/12,757) | 2.76%<br>(348/12,603) | 20.65%<br>(2,614/12,658) | 77.47%<br>(9,938/12,828) |
|  | B | 55.39%<br>(6,976/12,595) | 2.76%<br>(348/12,595) | 20.75%<br>(2,614/12,595) | 78.90%<br>(9,938/12,595) |
|  | C | 97.73%<br>(6976/7138) | 97.75%<br>(348/356) | 97.64%<br>(2614/2677) | 97.71%<br>(9938/10171) |
| *Caenorhabditis elegans* | A | 47.22%<br>(6,905/14,622) | 4.19%<br>(594/14,184) | 15.56%<br>(2,278/14,643) | 64.26%<br>(9,777/15,215) |
|  | B | 48.91%<br>(6,905/14,117) | 4.21%<br>(594/14,117) | 16.14%<br>(2,278/14,117) | 69.26%<br>(9,777/14,117) |
|  | C | 93.18%<br>(6,905/7,410) | 89.86%<br>(594/661) | 81.24%<br>(2,278/2,804) | 89.90%<br>(9,777/10,875) |
| *Danio rerio* | A | 59.28%<br>(8,389/14,152) | 3.77%<br>(533/14,152) | 27.68%<br>(3,917/14,152) | 90.72%<br>(12,839/14,152) |
|  | B | 59.28%<br>(8,389/14,152) | 3.77%<br>(533/14,152) | 27.68%<br>(3,917/14,152) | 90.72%<br>(12,839/14,152) |
|  | C | 100%<br>(8,389/8,389) | 100% (533/533) | 100%<br>(3,917/3,917) | 100%<br>(12,839/12,839) |
| *Mus musculus* | A | 63.21%<br>(7,392/11,695) | 5.73%<br>(659/11,495) | 17.70%<br>(2,173/12,280) | 81.26%<br>(10,224/12,582) |
|  | B | 64.59%<br>(7,392/11,444) | 5.76%<br>(659/11,444) | 18.99%<br>(2,173/11,444) | 89.34%<br>(10,224/11,444) |
|  | C | 96.72%<br>(7,392/7,643) | 92.82%<br>(659/710) | 72.22%<br>(2,173/3,009) | 89.98%<br>(10,224/11,362) |

The dynamics of all three indispensability parameters in the four studied species (*Drosophila melanogaster, Caenorhabditis elegans, Danio rerio* and *Mus musculus*) are presented in Figs. 2 and 3. In *D. melanogaster*, all three parameters show the same trends in phylogeny and ontogeny. The mean housekeeping potential (MHP) and the mean interaction potential (MIP) decreased, whereas *dN/dS* increased during phylogeny and ontogeny (Fig 2A, C, E; Suppl Figs. 1, 5, 9). The decrease of MIP was abrupt during the initial stages of both phylogeny and ontogeny, which resulted in the observed non-significance of the trends (Fig 2C; Suppl Fig 5). Together, these trends show that parallelism between phylogeny and ontogeny exists, and that recapitulation is observable from the beginning of ontogeny. In *C. elegans*, MHP and *dN/dS* showed recapitulation from both the beginning of ontogeny and from the phylotypic period (Fig 2B, F; Suppl Figs 2, 10), whereas MIP showed recapitulation from the beginning of ontogeny (Fig 2D; Suppl Fig 6). For two vertebrate species (*D. rerio* and *M. musculus*) recapitulation was observable from the phylotypic period for all parameters (Fig 3; Suppl Figs 3, 4, 7, 8, 11, 12). It is important to emphasize that our results are compatible with recapitulation after the phylotypic period, as demonstrated in all morphological examples of recapitulation in vertebrates (Swan 1990; Cohn and Tickle 1999; Lovejoy 2000; Metscher and Ahlberg 2001; Bejder and Hall 2002; Lovejoy *et al*. 2004; Thewissen *et al*. 2006; Nagashima *et al*. 2009; Cooper *et al*. 2014; Botelho *et al*. 2015; Diogo *et al*. 2015; Botelho *et al*. 2017; Thewissen 2018), as well as in the emergence of particular cis-regulatory elements in vertebrates (Uesaka et al. 2019).

**Figure 2.**
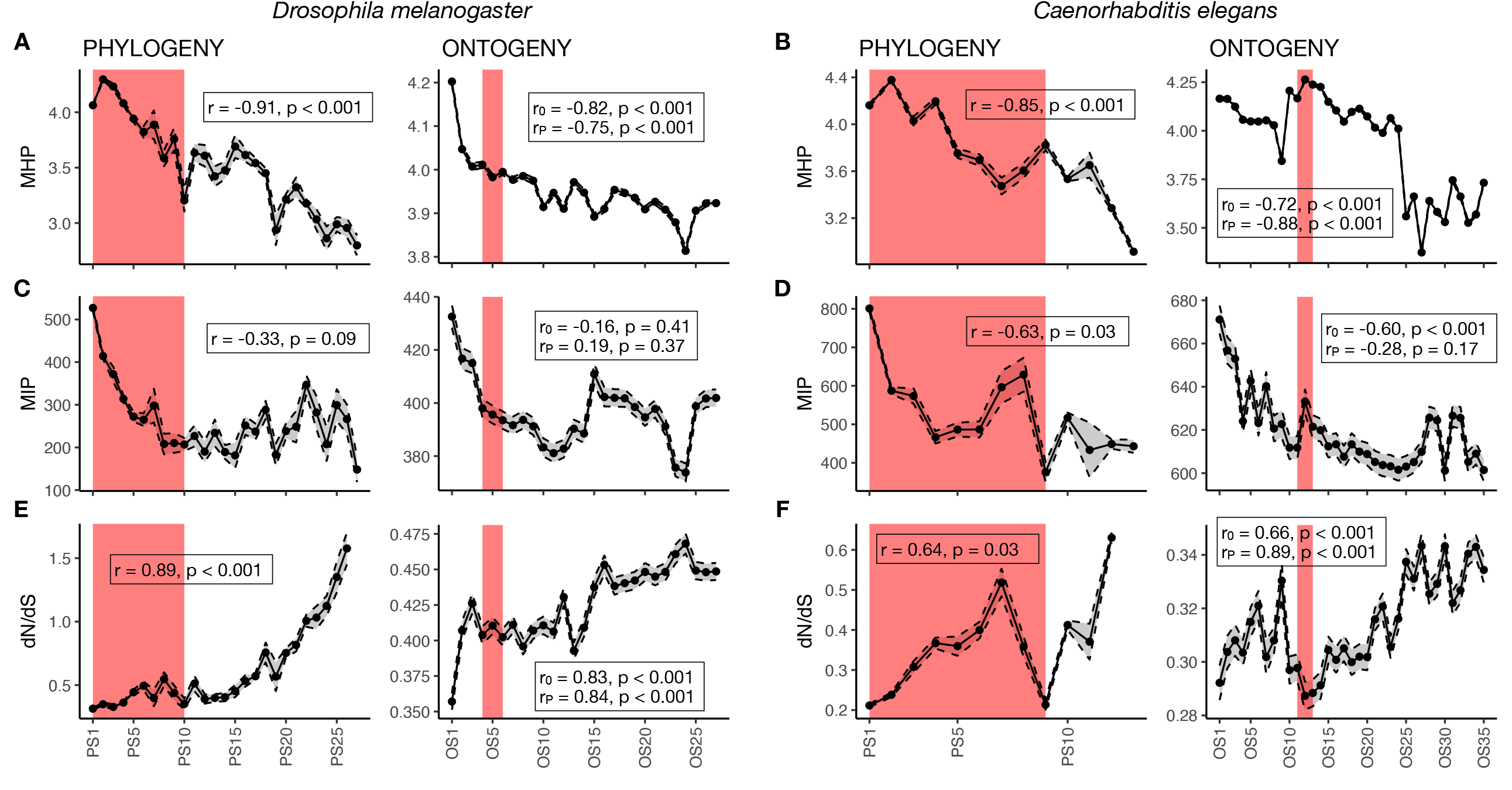
Parallelism between phylogeny and ontogeny. A, C, E. *Drosophila melanogaster*. B, D, F. *Caenorhabditis elegans*. A-B. Mean value (with associated standard error presented as a gray-shaded area) of housekeeping potential (MHP) for genes that belong to different phylostrata (PS; left panels), or are expressed in different ontogenetic stages (OS; right panels). C-D. Mean value (with associated standard error presented as a gray-shaded area) of the interaction potential (MIP) for genes that belong to different phylostrata (PS; left panels), or are expressed in different ontogenetic stages (OS; right panels). E-F. Mean value (with associated standard error presented as a gray-shaded area) of the *dN/dS* ratio for genes that belong to different phylostrata (PS; left panels), or are expressed in different ontogenetic stages (OS; right panels). The number associated with each phlyostratum (or ontogenetic stage) corresponds to their rank, with lower values signifying older phlyostrata (or earlier ontogenetic stages) and higher values signifying younger phlyostrata (or later ontogenetic stages). The pink-shaded areas correspond to the ontotypic period of phylogeny (left panels) and the phylotypic period of ontogeny (right panels), respectively. Spearman’s rank correlation coefficients (*r*) with associated *p*-values are reported for the relationship between the ranks of phylostrata (or ontogenetic stages) and A-B. MHP, C-D. MIP, E-F. *dN/dS*. In the case of ontogeny, the correlation was calculated either from the beginning of ontogeny *(r*_*0*_ coefficient) or the beginning of the phylotypic period (*r*_*P*_ coefficient). Data corresponds to the 0.1 expression quantile for the cut-off value.

**Figure 3.**
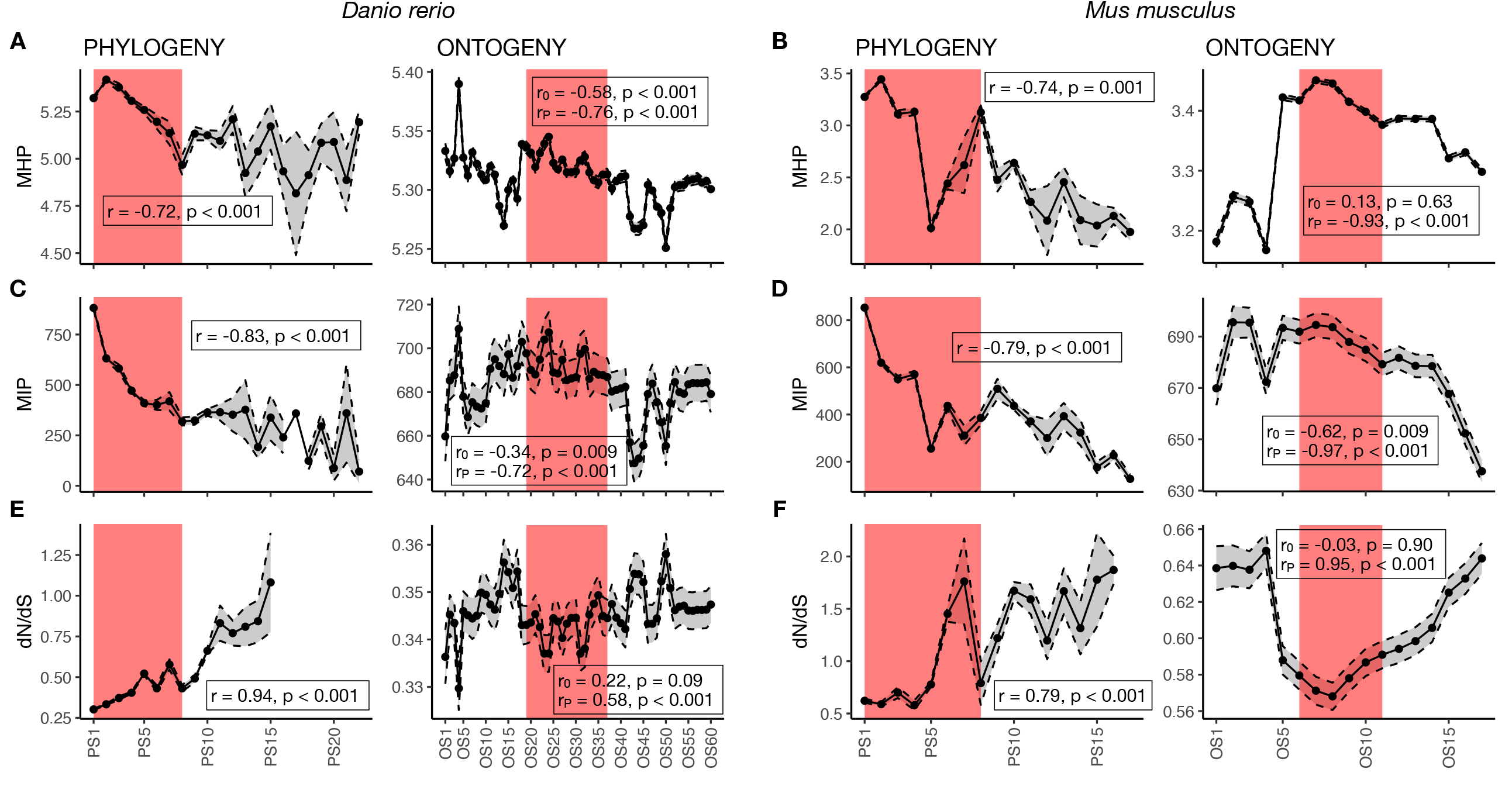
Parallelism between phylogeny and ontogeny. A, C, E. *Danio rerio*. B, D, F. *Mus musculus*. See Fig. 2 legend for details.

### Ontotypic period of phylogeny

Since parallelism between phylogeny and ontogeny exists, the next goal is to find the period in phylogeny which is analogous to the phylotypic period. This is important in order to know whether recapitulation starts from the beginning of ontogeny or from the phylotypic period. The phylotypic period is a period in ontogeny when genes responsible for the general body plan for the phylum are functionally expressed. Therefore, we wanted to find a period in phylogeny when these genes are formed. To answer this question we first used morphological criteria, and divided early ontogeny into three periods. These are Blastula (zygote, cleavage and formation of blastula), Gastrula (gastrulation and formation of gastrula) and the phylotypic period (general body plan for all species within a phylum). The period of phylogeny which corresponds to the phylotypic period we named the ontotypic period. The analogous period of Gastrula in phylogeny was named Gastrea, after Haeckel’s Gastrea theory (Haeckel 1872), whereas the analog to Blastula was named Blastea. In Blastea, we included phylostrata 1 (cellular organisms) and 2 (Eukaryota). Gastrea was equated to phylostratum 3 (Opisthokonta), since opisthoconts are ancestors of all Metazoa. In the ontotypic period, we included phylostratum 4 (Metazoa) and all subsequent phylostrata to the appropriate phylum, depending on the species.

To test whether the morphologically proposed analogous periods of phylogeny and ontogeny are biologically relevant, we compared the phylotypic period with three periods of phylogeny (Blastea, Gastrea and ontotypic period) based on their genic similarities. Since both ontogenetic and phylogenetic periods contain specific sets of genes, we defined genic similarity between two sets as the ratio of the number of elements (genes) in the intersection of two sets, to the number of genes in the union of the same two sets, i. e., 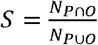, where *N*_*P*⋂*O*_ is the number of genes in the intersection of sets *P* (set of genes that emerged in specific phylogenetic periods) and *O* (set of genes that are functionally expressed in specific ontogenetic stages), and *N*_*P*⋃*O*_ is the number of genes in the union of these sets. *S* can take on any value from 0 to 1. The genic similarity of the phylotypic period was the highest with the Blastea period (*D. melanogaster* ∼ 0.55; *C. elegans* ∼ 0.47; *D. rerio* ∼ 0.59; *M. musculus* ∼ 0.63), followed with the ontotypic period (*D. melanogaster* ∼ 0.21; *C. elegans* ∼ 0.16; *D. rerio* ∼ 0.28; *M. musculus* ∼ 0.18), and the lowest similarity was observed with the Gastrea period (*D. melanogaster* ∼ 0.028; *C. elegans* ∼ 0.042; *D. rerio* ∼ 0.038; *M. musculus* ∼ 0.057) (Table 1; proportion A). We also analyzed the proportion of genes expressed in the phylotypic period with respect to their formation in phylogeny. In *D. melanogaster* ∼ 55 % of genes expressed in the phylotypic period emerged in Blastea, ∼ 21 % in the ontotypic period, whereas only ∼ 3 % emerged in Gastrea (Table 1; proportion B). For other species, the results are the following: *C. elegans* ∼ 49 % (Blastea), ∼ 16 % (ontotypic period), ∼ 4.2 % (Gastrea); *D. rerio* ∼ 59 % (Blastea), ∼ 28 % (ontotypic period), ∼ 3.8 % (Gastrea); *M. musculus* ∼ 64 % (Blastea), ∼ 19 % (ontotypic period), ∼ 5.8 % (Gastrea) (Table 1; proportion B).

From these results it is obvious that Blastea should be included in a period of phylogeny which is analogous to the phylotypic period of ontogeny. It could be included together with the ontotypic period or with both the ontotypic and Gastrea periods. Although Gastrea has a very small genic similarity with the phylotypic period, this is due to a large difference in the number of genes between the two sets. In order to decide whether to include or exclude Gastrea, we wanted to see which proportion of Gastrea genes is expressed in the phylotypic period. This can be estimated by the formula 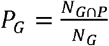, where *N*_*G*⋂*P*_ is the number of genes which are present in both Gastrea and the phylotypic period, and *N*_*G*_ is the total number of Gastrea genes. We found that almost all genes that emerged in Gastrea are expressed in the phylotypic period (Table 1; proportion C). Similar results were obtained with Blastea and the ontotypic period, as well as, with Blastea, Gastrea and ontotypic periods combined (BGO) (Table 1; proportion C). Therefore, all three morphologically defined periods should be included in a genetically defined ontotypic period. This is also supported by genic similarity of the phylotypic period with BGO (genetically defined ontotypic period). For small cut-offs, the genic similarities are ∼ 0.77 (*D. melanogaster*), ∼ 0.64 (*C. elegans*), ∼ 0.91 (*D. rerio*), and ∼ 0.81 (*M. musculus*) (Table 1; proportion A; Suppl. Table 3). Therefore, the genetically defined ontotypic period starts from the beginning of the phylogeny, and it is biologically relevant to consider that recapitulation starts from the phylotypic period (Figs. 1, 2 and 3).

### Recapitulation – a causal link between phylogeny and ontogeny

The last question we considered is the explanation for the origin of recapitulation, or the causal link between phylogeny and ontogeny. Haeckel assumed that ontogeny is caused by phylogeny (Haeckel 1866), whereas Muller had the opposite opinion (Muller 1864). At first glance, it seems that Muller was right since the phenomena at the population level of organization (phylogeny) are caused by the phenomena at the organismal level (ontogeny). However, for the explanation of parallelism between phylogeny and ontogeny this does not hold. The logic is the following: a novel gene is formed in the phylogeny, and after its formation it is expressed during ontogeny. Therefore, the cause occurs in the phylogeny, and the consequence is visible in ontogeny. From genomic and transcriptomic data, it follows that each gene is formed in one particular period in phylogeny, but it is usually expressed in several different periods in ontogeny. This implies that recapitulation (but not repetition) is possible. The critical condition to achieve recapitulation is that a gene which emerges later in phylogeny has higher chances to be expressed later in ontogeny. If this condition is satisfied we expect that, on average, phylogenetically younger genes will be expressed during later stages of ontogeny. Our results support this expectation (Fig 4; Suppl. Figs 13-16).

**Figure 4.**
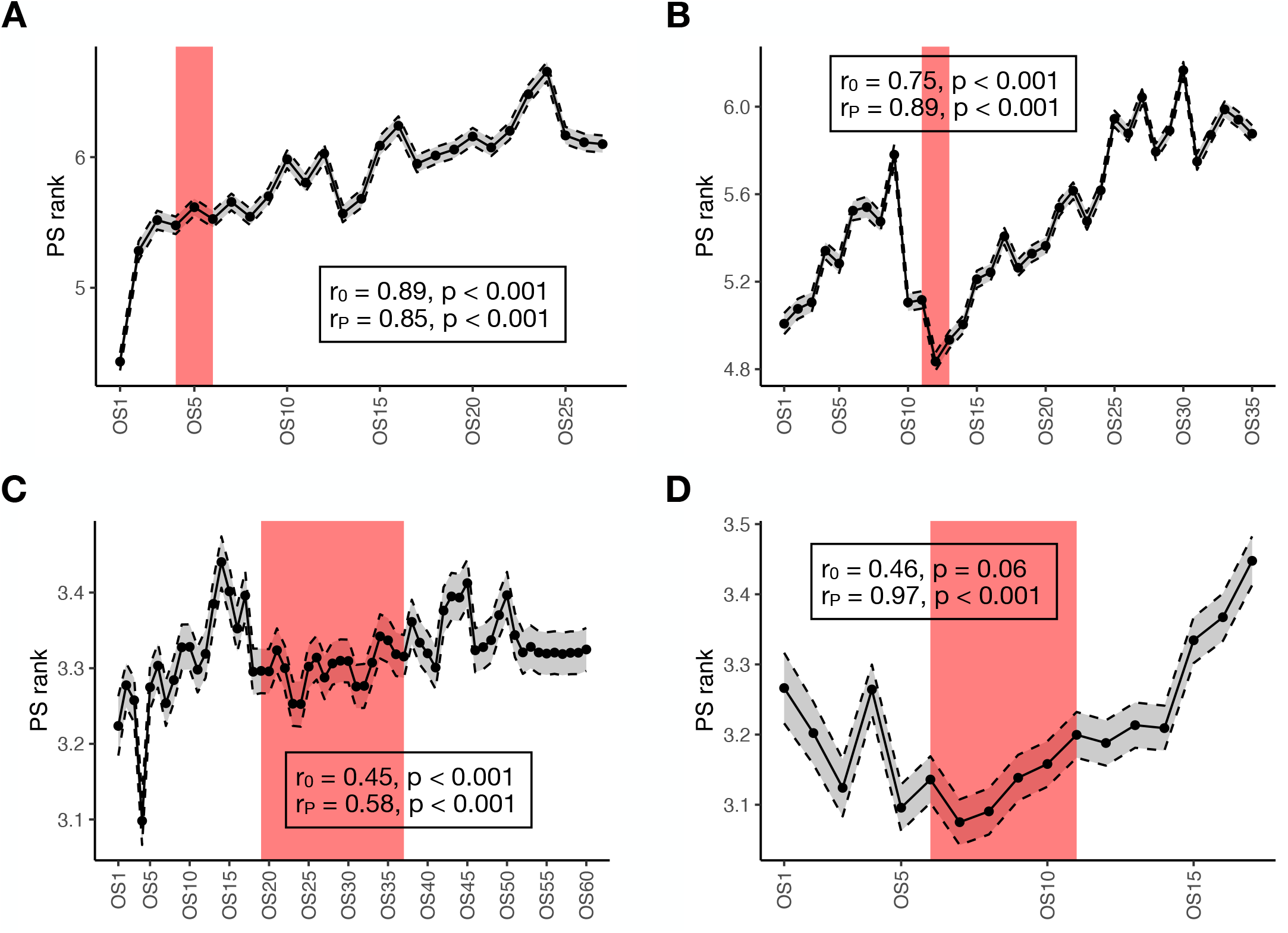
Mean phylogenetic age of genes expressed in different ontogenetic stages for A. *Drosophila melanogaster*, B. *Caenorhabditis elegans*, C. *Danio rerio* and D. *Mus musculus*. Mean phylogenetic age (with associated standard error presented as a gray-shaded area) is calculated as the mean phylostratum rank (PS rank) of genes expressed in different ontogenetic stages (OS). The number associated with each ontogenetic stage corresponds to its rank, with lower values signifying earlier ontogenetic stages and higher values signifying later ontogenetic stages. The pink-shaded area corresponds to the phylotypic period of ontogeny. The correlation between the mean phylostratum rank and the rank of ontogenetic stages was calculated either from the beginning of ontogeny (*r*_*0*_ coefficient) or the beginning of the phylotypic period (*r*_*P*_ coefficient). Data corresponds to the 0.1 expression quantile for the cut-off value.

## DISCUSSION

Although it is considered a modern version of the biogenetic law, the hourglass model does not provide any evidence for the existence of parallelism between phylogeny and ontogeny, or that ontogeny recapitulates phylogeny. Since the hourglass model and Haeckel’s law describe different phenomena, they are not mutually exclusive. In agreement with this, there are several morphological arguments in favour of recapitulation which are based on individual examples (Swan 1990; Cohn and Tickle 1999; Lovejoy 2000; Metscher and Ahlberg 2001; Bejder and Hall 2002; Lovejoy *et al*. 2004; Thewissen *et al*. 2006; Nagashima *et al*. 2009; Cooper *et al*. 2014; Botelho *et al*. 2015; Diogo *et al*. 2015; Botelho *et al*. 2017; Thewissen 2018), and one molecular example connected with chromatin accessibility (Uesaka *et al*. 2019). In contrast, there are morphological (Prum 1999; Bininda-Emonds *et al*. 2003; Poe 2006; Weisbecker *et al*. 2008; Richardson *et al*. 2009; Koyabu and Son 2014; Ziermann and Diogo 2014), as well as transcriptomic examples (Wang *et al*. 2013; Hu *et al*. 2017) for which attempts to demonstrate recapitulation failed.

We demonstrated that parallelism between phylogeny and ontogeny exists as a general phenomenon applicable to all animals. The mean indispensability of genes formed in the periods of phylogeny from the onset onwards decreases, as well as the mean indispensability of genes functionally expressed in the periods of ontogeny from the onset (or phylotypic period) onwards. Based on gene similarity, we defined the ontotypic period of phylogeny as the analogous period of the phylotypic period of ontogeny. The phylotypic period is a period of ontogeny when genes responsible for the resemblance between embryos of different species within a phylum are functionally expressed. On the other hand, the ontotypic period is the period of phylogeny when genes responsible for the similarity between embryos of different species within a phylum are formed. The ontotypic period contains genes that emerged at the beginning of phylogeny (phylostratum 1 related to cellular organisms) up to the phylostratum related to a particular phylum (Fig. 1). This observation is in favour of recapitulation starting from the phylotypic period of ontogeny. Before the phylotypic period, there are the periods characterized by species-specific features (shapes and sizes of cells, types of cell division and gastrulation) which develop into the evolutionary ancient and the most conserved (general) stage attributable to all species within a phylum (phylotypic stage). Parallelism, or recapitulation, is possible since genes which emerged in later periods of phylogeny have tendencies to be expressed in later periods of ontogeny (Fig. 4). If the periods of phylogeny and ontogeny from the beginning onwards share the similar genes, it could also be manifested at the phenotypic level, as morphological recapitulation as supported by the above-mentioned examples. In addition to parallelism, the correlation between genotype and phenotype is also visible in the conservation of morphological, as well as genomic and transcriptomic features during the phylotypic period of development (Hazkani Covo *et al*. 2005; Irie and Sehara-Fujisawa 2007; Kalinka et al. 2010; Irie and Kuratani 2011). Our results also showed that different species can follow different models of development. Vertebrates like mouse and zebrafish strongly follow the hourglass model with the highest average gene indispensability during the phylotypic period. On the other hand, *C. elegans* partially follows the hourglass model, whereas *D. melanogaster* follows the early conservation model of development.

The biogenetic law as formulated by Haeckel where "Ontogeny is a short and fast recapitulation of phylogeny” is based on comparative morphology and embryology, and is a thesis rather than a biological law based on quantitative regularities. Our results provide the quantitative and empirical regularities required to transform this thesis into a real biological law. Therefore, we reformulate the biogenetic law with the statement "Genes enter into the genetic program of ontogeny in a sequence similar to their emergence in phylogeny”.

## MATERIALS AND METHODS

### Data

Gene expression data for *Danio rerio, Drosophila melanogaster, Mus musculus* and *Caenorhabditis elegans* were taken from Domazet-Lošo and Tautz (2010), Graveley *et al*. (2011), Hu *et al*. (2017) and Li *et al*. (2014), respectively. To assess gene indispensability, we assessed three parameters associated with gene function (the housekeeping potential, the interaction potential and the *dN/dS* ratio, on a per-gene basis). We calculated the entropy of a gene’s expression (*H*_*g*_) across ontogenetic stages or phylostrata as described in Schug *et al*. (2005). We equate *H*_*g*_ with the housekeeping potential of a gene, where low *H*_*g*_ values indicate a high expression specificity of a gene, usually for a specific stage (and therefore low indispensability or low housekeeping potential), whereas high *H*_*g*_ values indicate a wide expression generality of a gene for many stages (and therefore high indispensability or high housekeeping potential). The mean housekeeping potential (MHP) was calculated as the mean value of housekeeping potentials for genes expressed in a particular ontogenetic period, or for genes that emerged in a particular phylostratum. Interaction data for each gene were taken from the STRING database v12.0 (Szklarczyk *et al*., 2023). Interaction potential was estimated for each (focal) gene as the number of genes that interact with the focal gene, as estimated by the STRING database. The mean interaction potential (MIP) was calculated as the mean value of interaction potentials for genes expressed in a particular ontogenetic period, or for genes that emerged in a particular phylostratum.

To asses per-gene *dN* and *dS* values we used the following reference sequences for the species: Release 6 (GenBank assembly GCA_000001215.4) for *D. melanogaster*, WS220 (downloaded from https://genome.ucsc.edu/) for *C. elegans*, GRCz10 (GenBank assembly GCA_000002035.3) for *D. rerio* and GRCm39 (GenBank assembly GCA_000001635.9) for *M. musculus*, downloaded from https://www.ncbi.nlm.nih.gov/. To calculate per-gene *dN* and *dS* values, we used the outgroups *Drosophila simulans* (GenBank assembly GCA_016746395.2), *Caenorhabditis brenneri* (GenBank assembly GCA_000143925.2), *Oryzias latipes* (GenBank assembly GCA_000313675.1) and *Mus spretus* (GenBank assembly GCA_921997135.2), for *D. melanogaster, C. elegans, D. rerio* and *M. musculus*, respectively. To determine the alignment of coding regions of references to their respective outgroup sequences, we used the liftOver program (Kuhn et al., 2013) with the corresponding UCSC chain files that contain the mapped genomic positions between the reference and the outgroup genome, downloaded from https://genome.ucsc.edu/. For all reference species, the *dN* (or *dS*) values were calculated on a per-gene basis as the ratio of codons that contain a non-synonymous (or synonymous) mutation with respect to the outgroup species, over the total number of alignable codons between the reference and outgroup gene. We only considered genes with at least 100 alignable nucleotides. The coding regions of each gene were determined using the GFF annotation files of the respective reference genomes. In figures, we show the mean *dN/dS* value for genes expressed in a particular ontogenetic period, or for genes that emerged in a particular phylostratum.

### Phylostratigraphy

Phylostratigraphy analysis was performed using the phylostratr R package (Arendsee et al., 2019) with default settings. Specifically, phylostratr retrieves a phylogenetic tree of species based on the NCBI tree of life (https://www.ncbi.nlm.nih.gov/), which is then trimmed such that the evolutionary distance between the remaining species is maximized. Proteomes of species were then downloaded from the UniProt database (https://www.uniprot.org/), followed by database construction and pairwise BLAST (Altschul et al., 1990) of the focal proteome against the proteome of each species in the trimmed phylogenetic tree. The quality of alignment between all query and target genes was based on the E-value (the expected number of random query-target alignments with an identical or better alignment score). The E-values were recorded and retained if above a predetermined cut-off (E-score < 10^−3^) signifying a presence of a gene within a phylogenetic clade. A phylostratum was then assigned to each gene as a ranked score defining the oldest phylogenetic clade containing the gene.

## Supporting information

Transcriptome Age Index

## ACKNOWLEDGMENTS

We thank Mary Sopta for improving the writing style of the manuscript.

## Author contributions

KBK conceived and conceptualized the work, and wrote the paper (main text). JB performed the bioinformatic analysis and wrote the paper (Materials and Methods section). RB contributed to bioinformatic analysis. All authors corrected and improved the first version of the manuscript.

## Competing interests

Authors have no competing interests to declare.

## Supplementary Figure legends

**Supplementary Figure 1**. Mean value (with associated standard error presented as a gray-shaded area) of housekeeping potential (MHP) for genes that are expressed in different ontogenetic stages (OS) for *Drosophila melanogaster* and given different expression cut-off values: A. Expression larger than 0. B. Expression larger than minimum gene expression. C-K. Expression larger than the specified quantile of gene expression. L. Expression equal to maximum gene expression. The number associated with each ontogenetic stage corresponds to its rank, with lower values signifying earlier ontogenetic stages and higher values signifying later ontogenetic stages. The pink-shaded area corresponds to the phylotypic period of ontogeny. The correlation between MHP and the rank of ontogenetic stages was calculated either from the beginning of ontogeny (*r*_*0*_ coefficient) or the beginning of the phylotypic period (*r*_*P*_ coefficient).

**Supplementary Figure 2**. Mean value (with associated standard error presented as a gray-shaded area) of housekeeping potential (MHP) for genes that are expressed in different ontogenetic stages (OS) for *Caenorhabditis elegans* and given different expression cut-off values: A. Expression larger than 0. B. Expression larger than minimum gene expression. C-K. Expression larger than the specified quantile of gene expression. L. Expression equal to maximum gene expression. The number associated with each ontogenetic stage corresponds to its rank, with lower values signifying earlier ontogenetic stages and higher values signifying later ontogenetic stages. The pink-shaded area corresponds to the phylotypic period of ontogeny. The correlation between MHP and the rank of ontogenetic stages was calculated either from the beginning of ontogeny (*r*_*0*_ coefficient) or the beginning of the phylotypic period (*r*_*P*_ coefficient).

**Supplementary Figure 3**. Mean value (with associated standard error presented as a gray-shaded area) of housekeeping potential (MHP) for genes that are expressed in different ontogenetic stages (OS) for *Danio rerio* and given different expression cut-off values: A. Expression larger than 0. B. Expression larger than minimum gene expression. C-K. Expression larger than the specified quantile of gene expression. L. Expression equal to maximum gene expression. The number associated with each ontogenetic stage corresponds to its rank, with lower values signifying earlier ontogenetic stages and higher values signifying later ontogenetic stages. The pink-shaded area corresponds to the phylotypic period of ontogeny. The correlation between MHP and the rank of ontogenetic stages was calculated either from the beginning of ontogeny (*r*_*0*_ coefficient) or the beginning of the phylotypic period (*r*_*P*_ coefficient).

**Supplementary Figure 4**. Mean value (with associated standard error presented as a gray-shaded area) of housekeeping potential (MHP) for genes that are expressed in different ontogenetic stages (OS) for *Mus musculus* and given different expression cut-off values: A. Expression larger than 0. B. Expression larger than minimum gene expression. C-K. Expression larger than the specified quantile of gene expression. L. Expression equal to maximum gene expression. The number associated with each ontogenetic stage corresponds to its rank, with lower values signifying earlier ontogenetic stages and higher values signifying later ontogenetic stages. The pink-shaded area corresponds to the phylotypic period of ontogeny. The correlation between MHP and the rank of ontogenetic stages was calculated either from the beginning of ontogeny (*r*_*0*_ coefficient) or the beginning of the phylotypic period (*r*_*P*_ coefficient).

**Supplementary Figure 5**. Mean value (with associated standard error presented as a gray-shaded area) of the interaction potential (MIP) for genes that are expressed in different ontogenetic stages (OS) for *Drosophila melanogaster* and given different expression cut-off values: A. Expression larger than 0. B. Expression larger than minimum gene expression. C-K. Expression larger than the specified quantile of gene expression. L. Expression equal to maximum gene expression. The number associated with each ontogenetic stage corresponds to its rank, with lower values signifying earlier ontogenetic stages and higher values signifying later ontogenetic stages. The pink-shaded area corresponds to the phylotypic period of ontogeny. The correlation between MIP and the rank of ontogenetic stages was calculated either from the beginning of ontogeny (*r*_*0*_ coefficient) or the beginning of the phylotypic period (*r*_*P*_ coefficient).

**Supplementary Figure 6**. Mean value (with associated standard error presented as a gray-shaded area) of the interaction potential (MIP) for genes that are expressed in different ontogenetic stages (OS) for *Caenorhabditis elegans* and given different expression cut-off values: A. Expression larger than 0. B. Expression larger than minimum gene expression. C-K. Expression larger than the specified quantile of gene expression. L. Expression equal to maximum gene expression. The number associated with each ontogenetic stage corresponds to its rank, with lower values signifying earlier ontogenetic stages and higher values signifying later ontogenetic stages. The pink-shaded area corresponds to the phylotypic period of ontogeny. The correlation between MIP and the rank of ontogenetic stages was calculated either from the beginning of ontogeny (*r*_*0*_ coefficient) or the beginning of the phylotypic period (*r*_*P*_ coefficient).

**Supplementary Figure 7**. Mean value (with associated standard error presented as a gray-shaded area) of the interaction potential (MIP) for genes that are expressed in different ontogenetic stages (OS) for *Danio rerio* and given different expression cut-off values: A. Expression larger than 0. B. Expression larger than minimum gene expression. C-K. Expression larger than the specified quantile of gene expression. L. Expression equal to maximum gene expression. The number associated with each ontogenetic stage corresponds to its rank, with lower values signifying earlier ontogenetic stages and higher values signifying later ontogenetic stages. The pink-shaded area corresponds to the phylotypic period of ontogeny. The correlation between MIP and the rank of ontogenetic stages was calculated either from the beginning of ontogeny (*r*_*0*_ coefficient) or the beginning of the phylotypic period (*r*_*P*_ coefficient).

**Supplementary Figure 8**. Mean value (with associated standard error presented as a gray-shaded area) of the interaction potential (MIP) for genes that are expressed in different ontogenetic stages (OS) for *Mus musculus* and given different expression cut-off values: A. Expression larger than 0. B. Expression larger than minimum gene expression. C-K. Expression larger than the specified quantile of gene expression. L. Expression equal to maximum gene expression. The number associated with each ontogenetic stage corresponds to its rank, with lower values signifying earlier ontogenetic stages and higher values signifying later ontogenetic stages. The pink-shaded area corresponds to the phylotypic period of ontogeny. The correlation between MIP and the rank of ontogenetic stages was calculated either from the beginning of ontogeny (*r*_*0*_ coefficient) or the beginning of the phylotypic period (*r*_*P*_ coefficient).

**Supplementary Figure 9**. Mean value (with associated standard error presented as a gray-shaded area) of the *dN/dS* ratio for genes that are expressed in different ontogenetic stages (OS) for *Drosophila melanogaster* and given different expression cut-off values: A. Expression larger than 0. B. Expression larger than minimum gene expression. C-K. Expression larger than the specified quantile of gene expression. L. Expression equal to maximum gene expression. The number associated with each ontogenetic stage corresponds to its rank, with lower values signifying earlier ontogenetic stages and higher values signifying later ontogenetic stages. The pink-shaded area corresponds to the phylotypic period of ontogeny. The correlation between *dN/dS* and the rank of ontogenetic stages was calculated either from the beginning of ontogeny (*r*_*0*_ coefficient) or the beginning of the phylotypic period (*r*_*P*_ coefficient).

**Supplementary Figure 10**. Mean value (with associated standard error presented as a gray-shaded area) of the *dN/dS* ratio for genes that are expressed in different ontogenetic stages (OS) for *Caenorhabditis elegans* and given different expression cut-off values: A. Expression larger than 0. B. Expression larger than minimum gene expression. C-K. Expression larger than the specified quantile of gene expression. L. Expression equal to maximum gene expression. The number associated with each ontogenetic stage corresponds to its rank, with lower values signifying earlier ontogenetic stages and higher values signifying later ontogenetic stages. The pink-shaded area corresponds to the phylotypic period of ontogeny. The correlation between *dN/dS* and the rank of ontogenetic stages was calculated either from the beginning of ontogeny (*r*_*0*_ coefficient) or the beginning of the phylotypic period (*r*_*P*_ coefficient).

**Supplementary Figure 11**. Mean value (with associated standard error presented as a gray-shaded area) of the *dN/dS* ratio for genes that are expressed in different ontogenetic stages (OS) for *Danio rerio* and given different expression cut-off values: A. Expression larger than 0. B. Expression larger than minimum gene expression. C-K. Expression larger than the specified quantile of gene expression. L. Expression equal to maximum gene expression. The number associated with each ontogenetic stage corresponds to its rank, with lower values signifying earlier ontogenetic stages and higher values signifying later ontogenetic stages. The pink-shaded area corresponds to the phylotypic period of ontogeny. The correlation between *dN/dS* and the rank of ontogenetic stages was calculated either from the beginning of ontogeny *(r*_*0*_ coefficient) or the beginning of the phylotypic period (*r*_*P*_ coefficient).

**Supplementary Figure 12**. Mean value (with associated standard error presented as a gray-shaded area) of the *dN/dS* ratio for genes that are expressed in different ontogenetic stages (OS) for *Mus musculus* and given different expression cut-off values: A. Expression larger than 0. B. Expression larger than minimum gene expression. C-K. Expression larger than the specified quantile of gene expression. L. Expression equal to maximum gene expression. The number associated with each ontogenetic stage corresponds to its rank, with lower values signifying earlier ontogenetic stages and higher values signifying later ontogenetic stages. The pink-shaded area corresponds to the phylotypic period of ontogeny. The correlation between *dN/dS* and the rank of ontogenetic stages was calculated either from the beginning of ontogeny (*r*_*0*_ coefficient) or the beginning of the phylotypic period *(r*_*P*_ coefficient).

**Supplementary Figure 13**. Mean phylogenetic age (with associated standard error presented as a gray-shaded area) of genes (PS rank) that are expressed in different ontogenetic stages (OS) for *Drosophila melanogaster* and given different expression cut-off values: A. Expression larger than 0. B. Expression larger than minimum gene expression. C-K. Expression larger than the specified quantile of gene expression. L. Expression equal to maximum gene expression. The number associated with each ontogenetic stage corresponds to its rank, with lower values signifying earlier ontogenetic stages and higher values signifying later ontogenetic stages. The pink-shaded area corresponds to the phylotypic period of ontogeny. The correlation between the PS rank and the rank of ontogenetic stages was calculated either from the beginning of ontogeny (*r*_*0*_ coefficient) or the beginning of the phylotypic period (*r*_*P*_ coefficient).

**Supplementary Figure 14**. Mean phylogenetic age (with associated standard error presented as a gray-shaded area) of genes (PS rank) that are expressed in different ontogenetic stages (OS) for *Caenorhabditis elegans* and given different expression cut-off values: A. Expression larger than 0. B. Expression larger than minimum gene expression. C-K. Expression larger than the specified quantile of gene expression. L. Expression equal to maximum gene expression. The number associated with each ontogenetic stage corresponds to its rank, with lower values signifying earlier ontogenetic stages and higher values signifying later ontogenetic stages. The pink-shaded area corresponds to the phylotypic period of ontogeny. The correlation between the PS rank and the rank of ontogenetic stages was calculated either from the beginning of ontogeny (*r*_*0*_ coefficient) or the beginning of the phylotypic period (*r*_*P*_ coefficient).

**Supplementary Figure 15**. Mean phylogenetic age (with associated standard error presented as a gray-shaded area) of genes (PS rank) that are expressed in different ontogenetic stages (OS) for *Danio rerio* and given different expression cut-off values: A. Expression larger than 0. B. Expression larger than minimum gene expression. C-K. Expression larger than the specified quantile of gene expression. L. Expression equal to maximum gene expression. The number associated with each ontogenetic stage corresponds to its rank, with lower values signifying earlier ontogenetic stages and higher values signifying later ontogenetic stages. The pink-shaded area corresponds to the phylotypic period of ontogeny. The correlation between the PS rank and the rank of ontogenetic stages was calculated either from the beginning of ontogeny (*r*_*0*_ coefficient) or the beginning of the phylotypic period (*r*_*P*_ coefficient).

**Supplementary Figure 16**. Mean phylogenetic age (with associated standard error presented as a gray-shaded area) of genes (PS rank) that are expressed in different ontogenetic stages (OS) for *Mus musculus* and given different expression cut-off values: A. Expression larger than 0. B. Expression larger than minimum gene expression. C-K. Expression larger than the specified quantile of gene expression. L. Expression equal to maximum gene expression. The number associated with each ontogenetic stage corresponds to its rank, with lower values signifying earlier ontogenetic stages and higher values signifying later ontogenetic stages. The pink-shaded area corresponds to the phylotypic period of ontogeny. The correlation between the PS rank and the rank of ontogenetic stages was calculated either from the beginning of ontogeny (*r*_*0*_ coefficient) or the beginning of the phylotypic period (*r*_*P*_ coefficient).

## Supplementary Table legends

Supplementary Table 1. Data associated to Figs. 2-4 and Supplementary Figs. 1-16.

Supplementary Table 2. Correlation coefficients from Figs. 2-4 and Supplementary Figs. 1-16.

Supplementary Table 3. Proportions from Table 1 for different expression cut-off value

**SUPPLEMENT 1**

**Transcriptome Age Index**

**SUPPLEMENT 2**

**Supplementary Figures**

**SUPPLEMENT 3**

**Supplementary Tables**

## Notes

### Competing Interest Statement

The authors have declared no competing interest.

