## Supplementary material for "Parallelism between phylogeny and ontogeny": Transcriptome Age Index

**SUPPLEMENT 1**

**Transcriptome Age Index**

Transcriptome Age Index ($TAI$) is not an adequate parameter to estimate the average age of genes expressed in transcriptome if the age and expression of genes are correlated. $TAI$ is defined by the formula

$$TAI=\frac{\sum_{i=1}^{n} A_{i}E_{i}}{\sum_{i=1}^{n} E_{i}}$$

where $A_{i}$ is the age of *i*-th gene (the number of its phylostratum), $E_{i}$ is its level of expression in particular stage of ontogeny, and *n* is the total number of genes expressed in this stage of ontogeny (Domazet-Lošo and Tautz 2010). The denominator of $TAI$ for particular stage of ontogeny is always the same, whereas numerator is biased by the level of expression if the age and level of expression of genes are correlated. Consequently, if older genes are much more expressed than younger, then result is smaller, i. e. the transcriptome seems older. In opposite situation, the result is larger, i. e. the transcriptome seems younger. If correlation exists, $TAI$ can produce a random deviation or artefact, from which it is not possible to make biologically relevant conclusion. The appropriate way to estimate the average age of functionally expressed genes in particular period of ontogeny is simply

$$\bar{A}=\frac{1}{n}\sum_{i=1}^{n} A_{i}$$

but in this case one should choose the cut-off value of expression level above which the genes are biologically functional. Usually, one should use several cut-off values to see if there is a regularity of trends. $TAI$ would be equal to $\bar{A}$ if the expression level of all genes are the same, or if there is no correlation between the age and the expression level of genes.

In the simplest condition, when the expression level is equal (constant) to each gene ($E_{i}=E$) we can write

$$TAI=\frac{E\sum_{i=1}^{n} A_{i}}{nE}=\frac{1}{n}\sum_{i=1}^{n} A_{i}=\bar{A}$$

and $TAI$ is equal to $\bar{A}$ under the single cut-off value (cut-off $E_{i}>0$). This means that each expressed gene is biologically functional.

The first criticism of $TAI$ came from Robinson-Rechavi group (Piasecka *et al.* 2013) who re-analysed data from paper (Domazet-Lošo and Tautz 2010). In order to reduce the effect of highly expressed genes, the original expression data were ${log}_{10}$-transformed, and the result was different, i. e., the hourglass model was transformed into early-conservation model. The difference between the least and highest expressed genes in Zebrafish is 5 order of magnitude which implies that $TAI$ could be sensitive to the effect of outliers. When they removed just one outlier with high expression the outcome was different, as well as the conclusion. On the other hand, the ${log}_{10}$-transformed data were resistant to the effect of outliers. This means that decrease in the expression difference between genes (decrease in difference of weighting factors for gene age in $TAI$ formula) can change the overall pattern of the age of transcriptome during development. They also analysed the age of transcriptome as the average age of genes expressed in particular stage of development using one cut-off value (cut-off ${log}_{10}E_{i}>1$). The result was similar with ${log}_{10}$-transformed data in $TAI$ formula. For more details of this analysis see the reference (Piasecka *et al*. 2013).

In conclusion about estimation of the age of transcriptome, we will emphasize three points: 1) the gene function is biologically relevant, but not the level of expression above which it is established; 2) it is not realistically to expect that there is no correlation between gene age and its expression level at any stage of development; and 3) even if there is no correlation between the gene age and its expression $TAI$ becomes $\bar{A}$ with the single cut-off value of expression (cut-off $E_{i}>0$), but for the adequate analysis one should use multiple cut-off values to see the trends made by functional genes. For example, in our analysis of mean indispensability of genes expressed in particular periods of ontogeny, we used 12 cut-off values in order to make relevant conclusion (Supplementary Figures). Therefore, the above-mentioned facts argue that $TAI$ is not suitable for estimation of the average age of genes expressed in transcriptome.
