## Supplementary figures and images for "Parallelism between phylogeny and ontogeny"

### supp_fig1.pdf

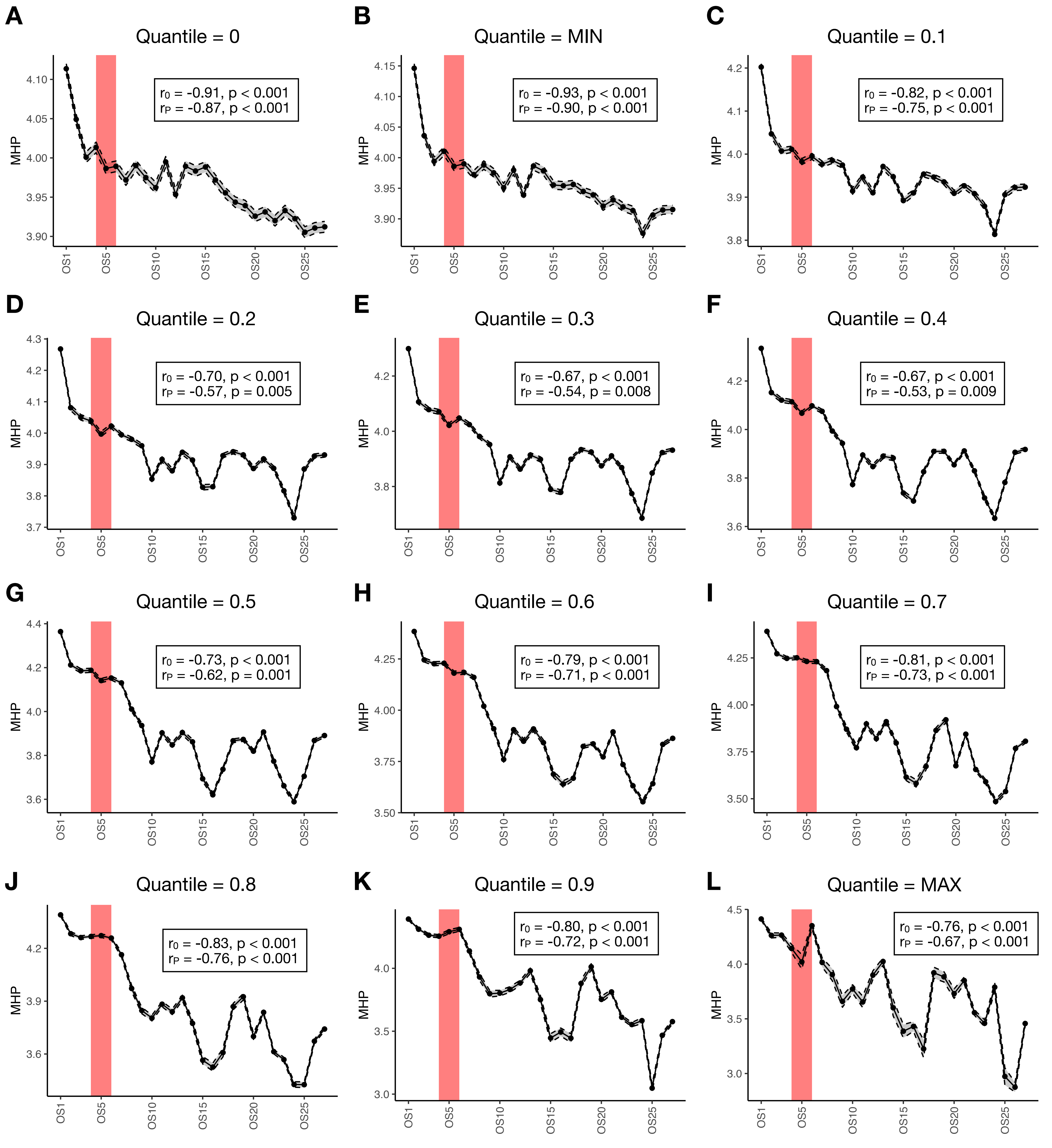

### supp_fig2.pdf

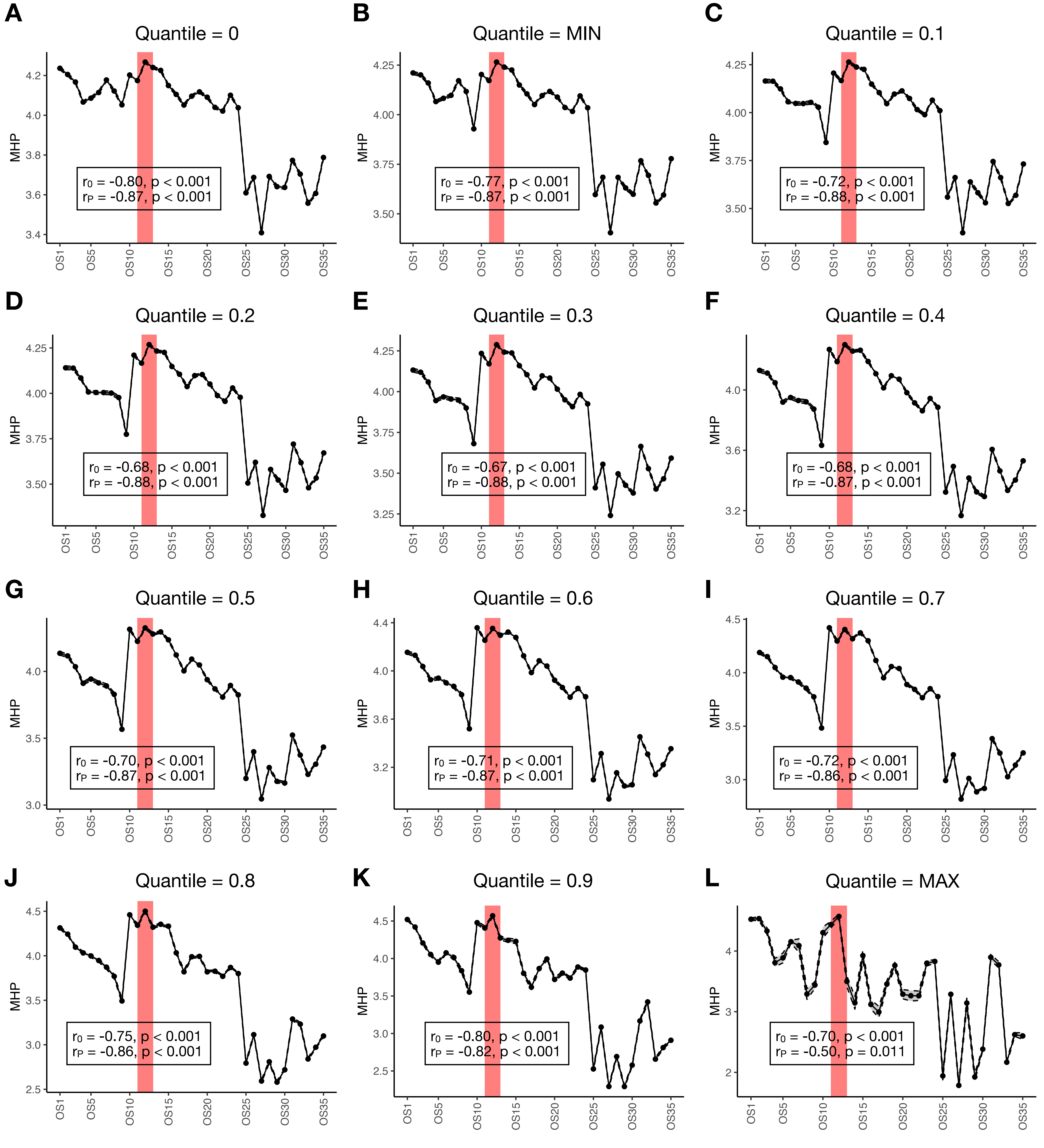

### supp_fig3.pdf

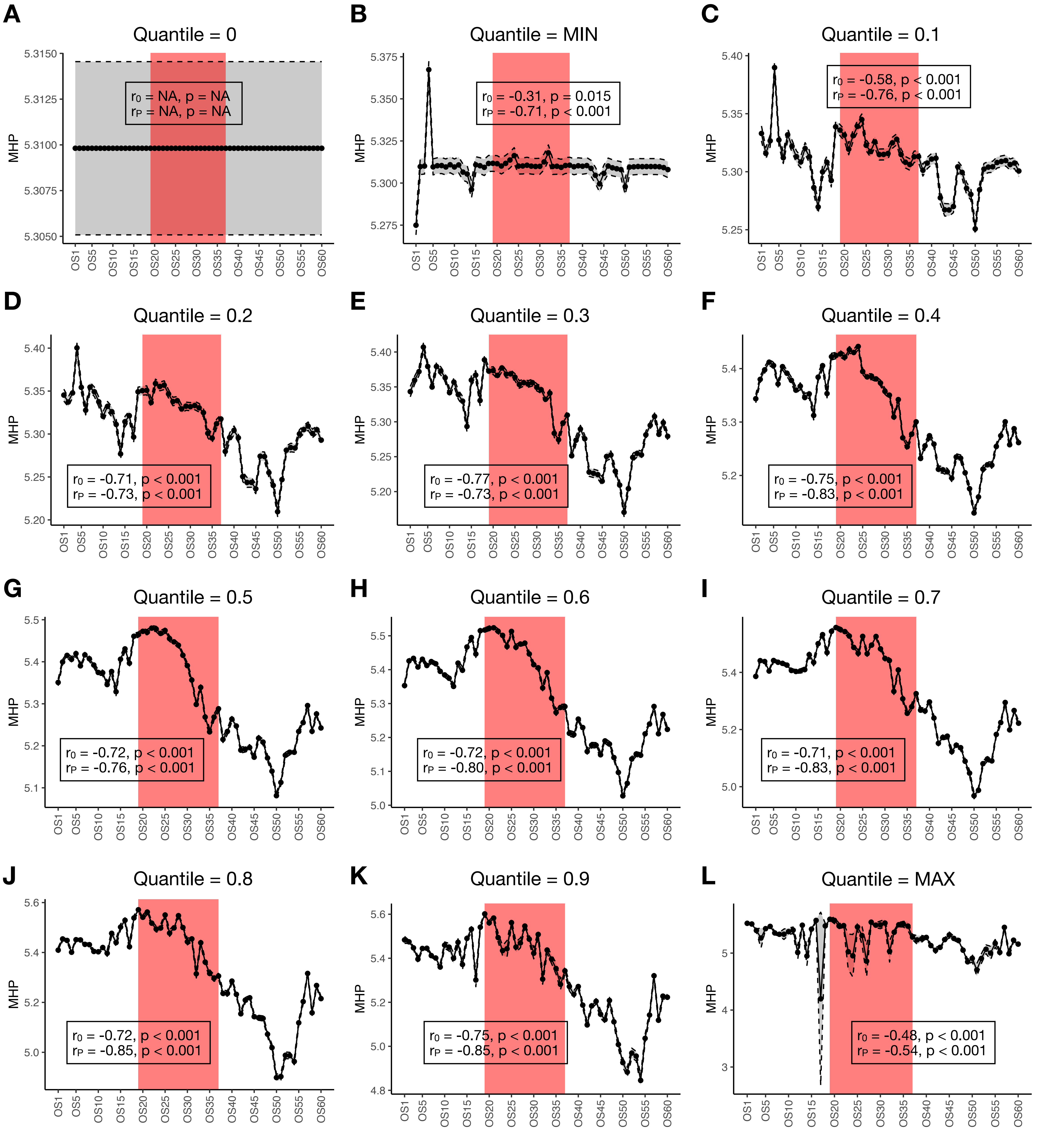

### supp_fig4.pdf

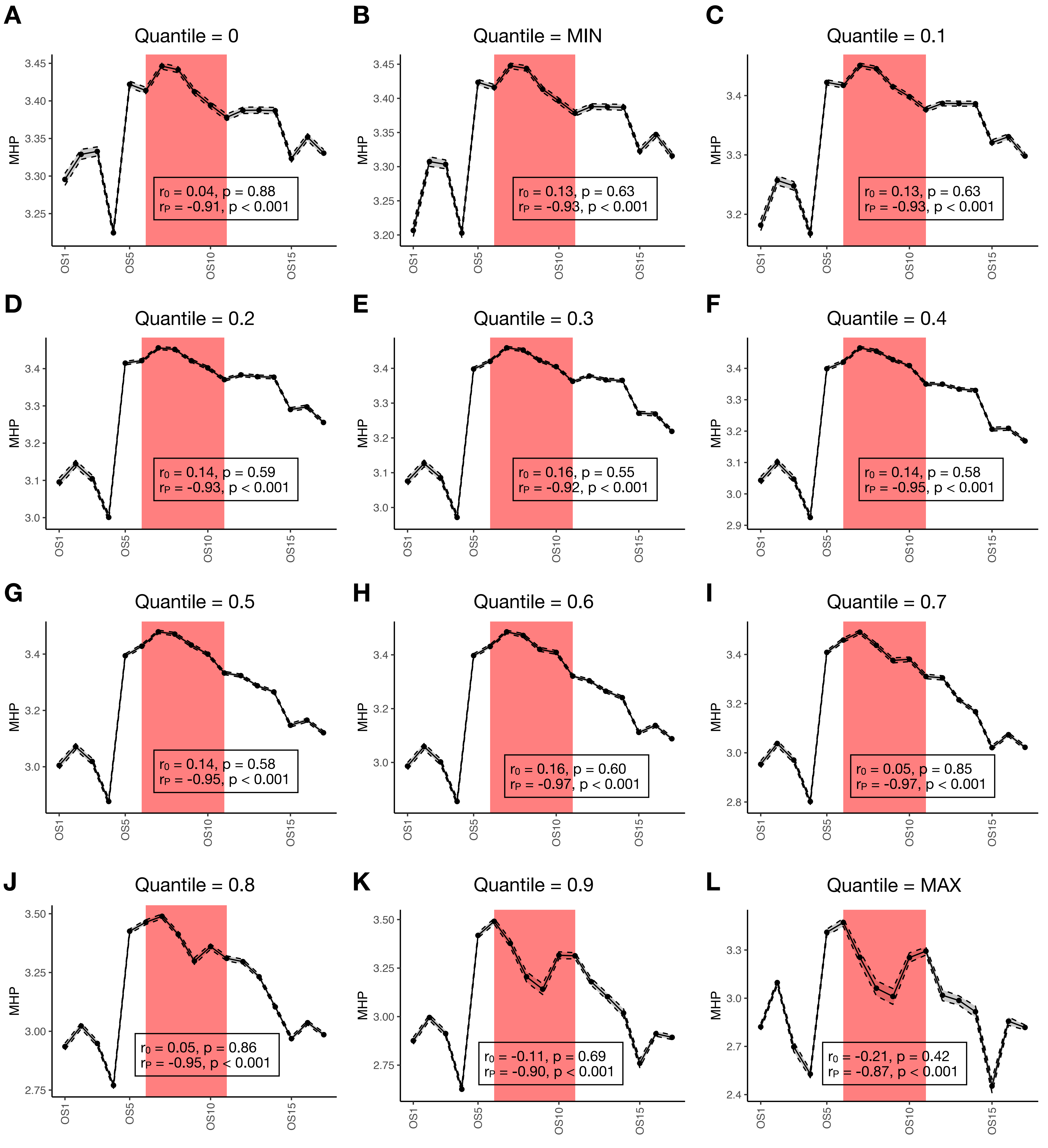

### supp_fig5.pdf

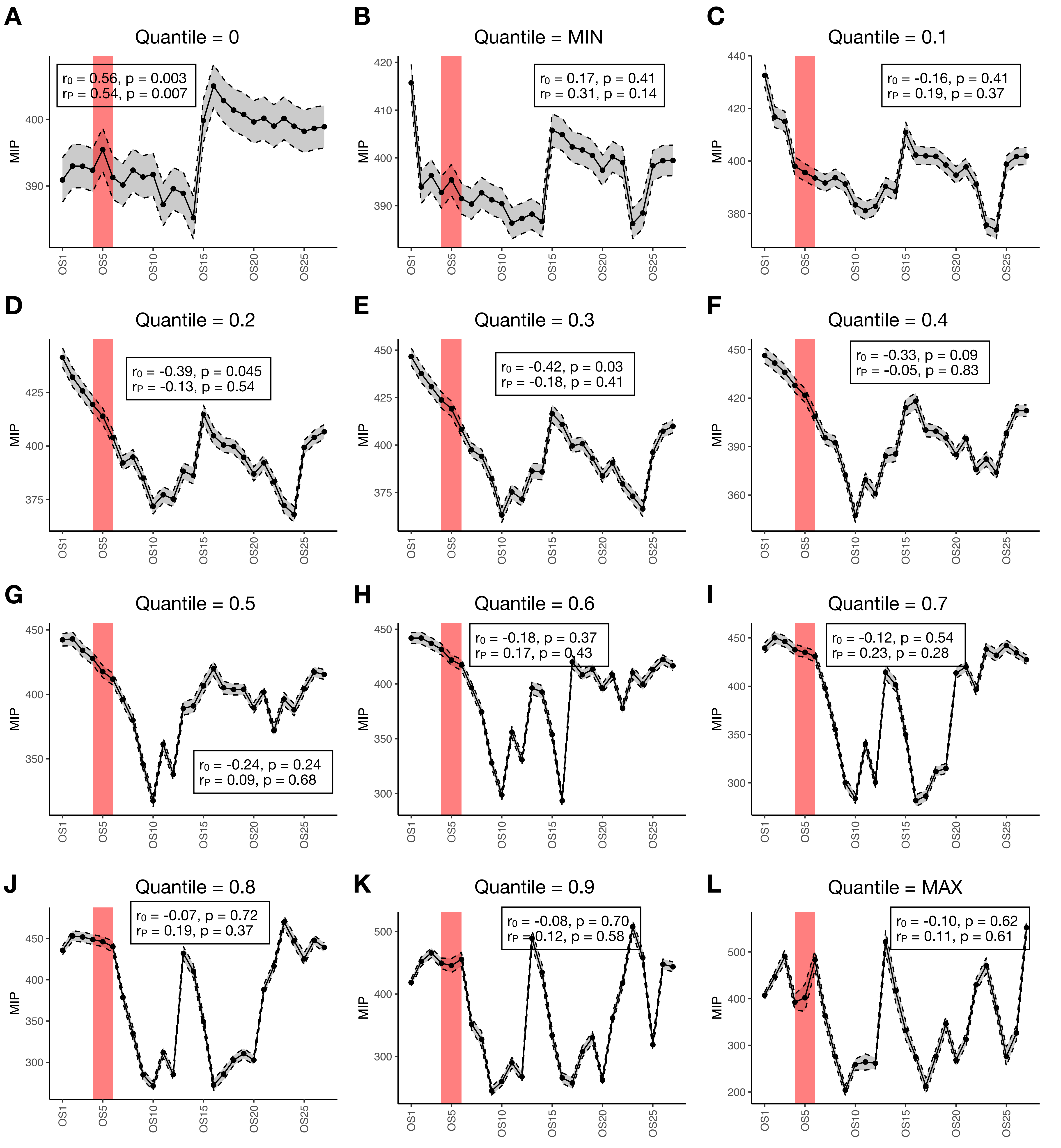

### supp_fig6.pdf

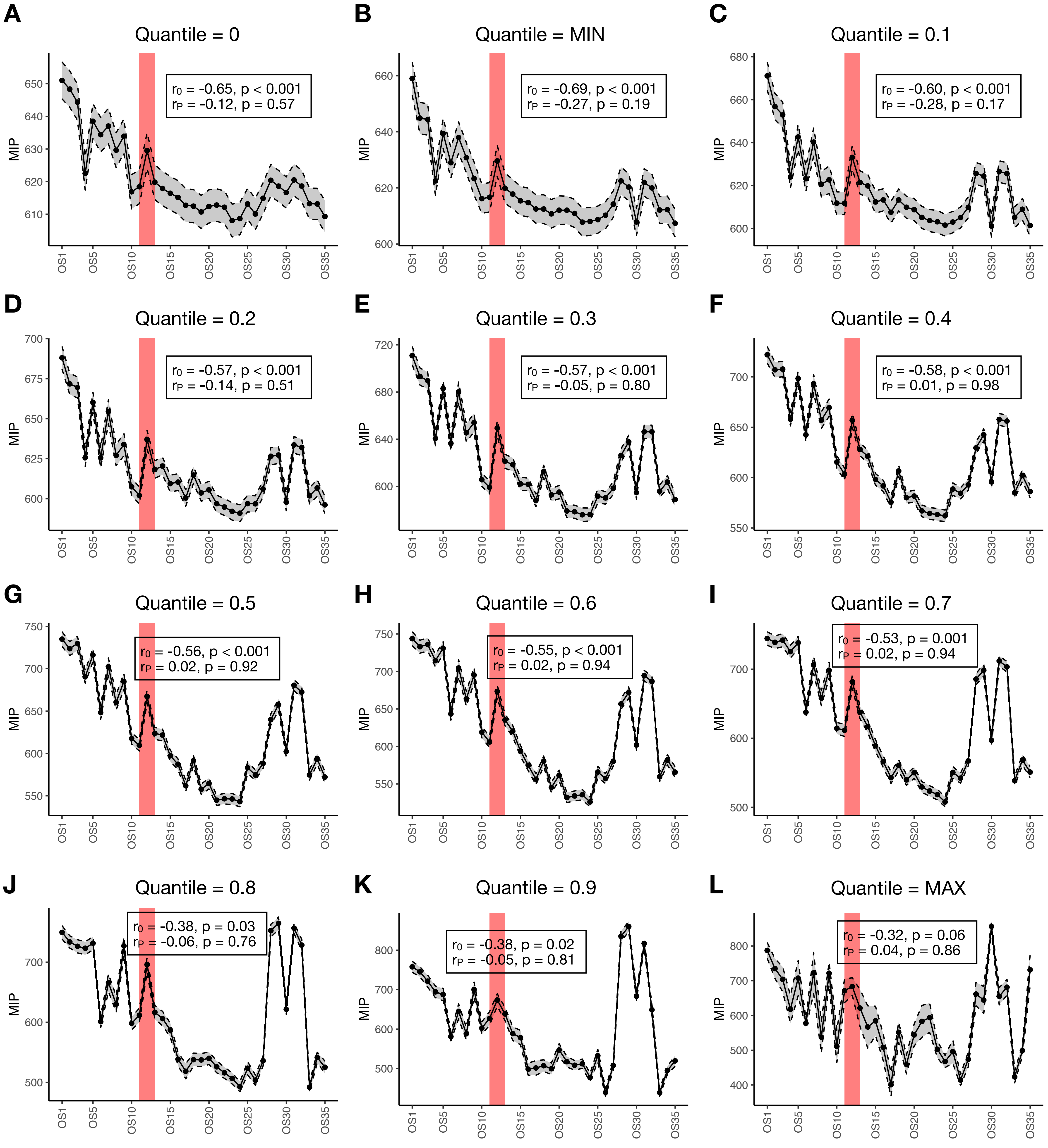

### supp_fig7.pdf

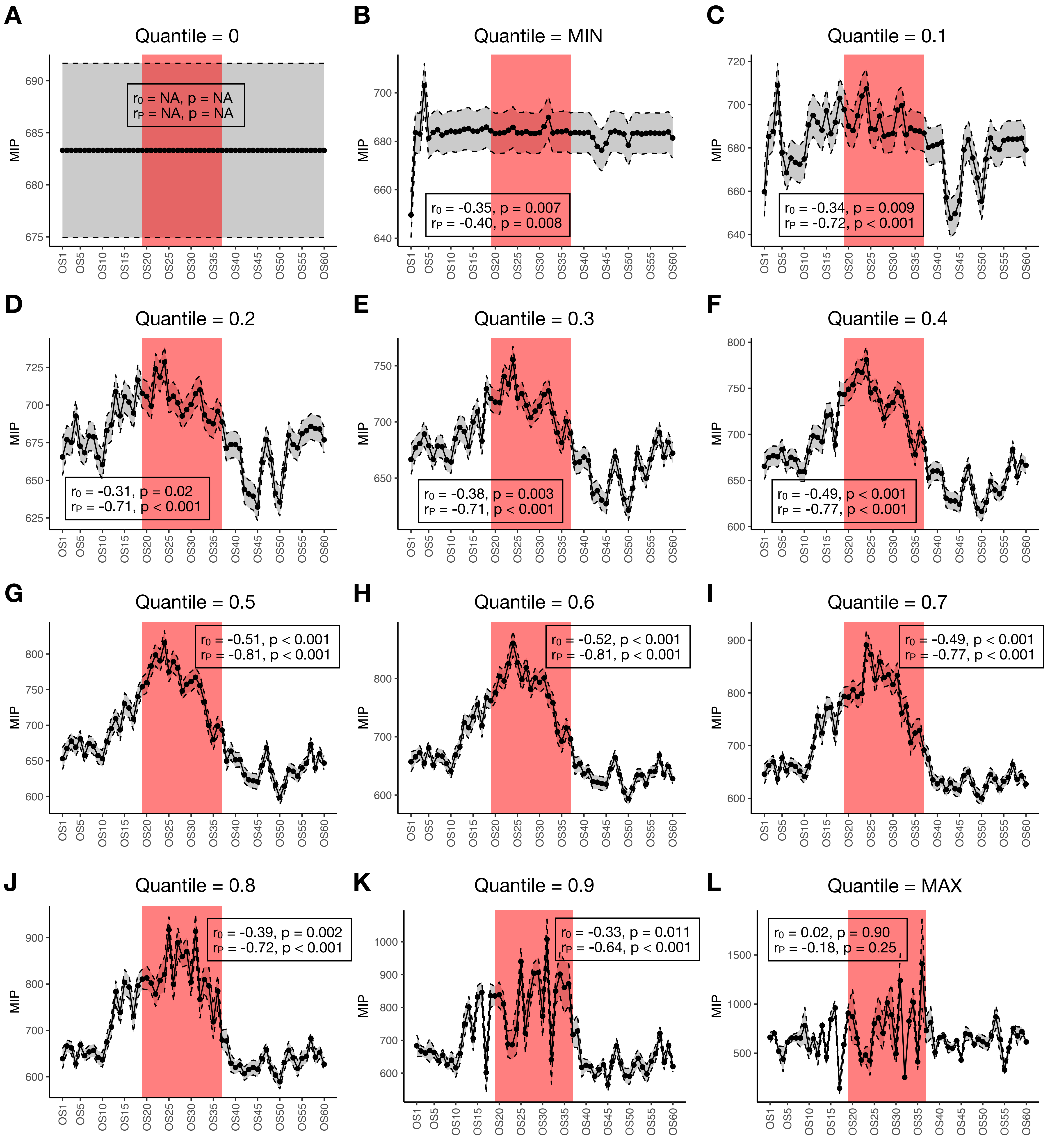

### supp_fig8.pdf

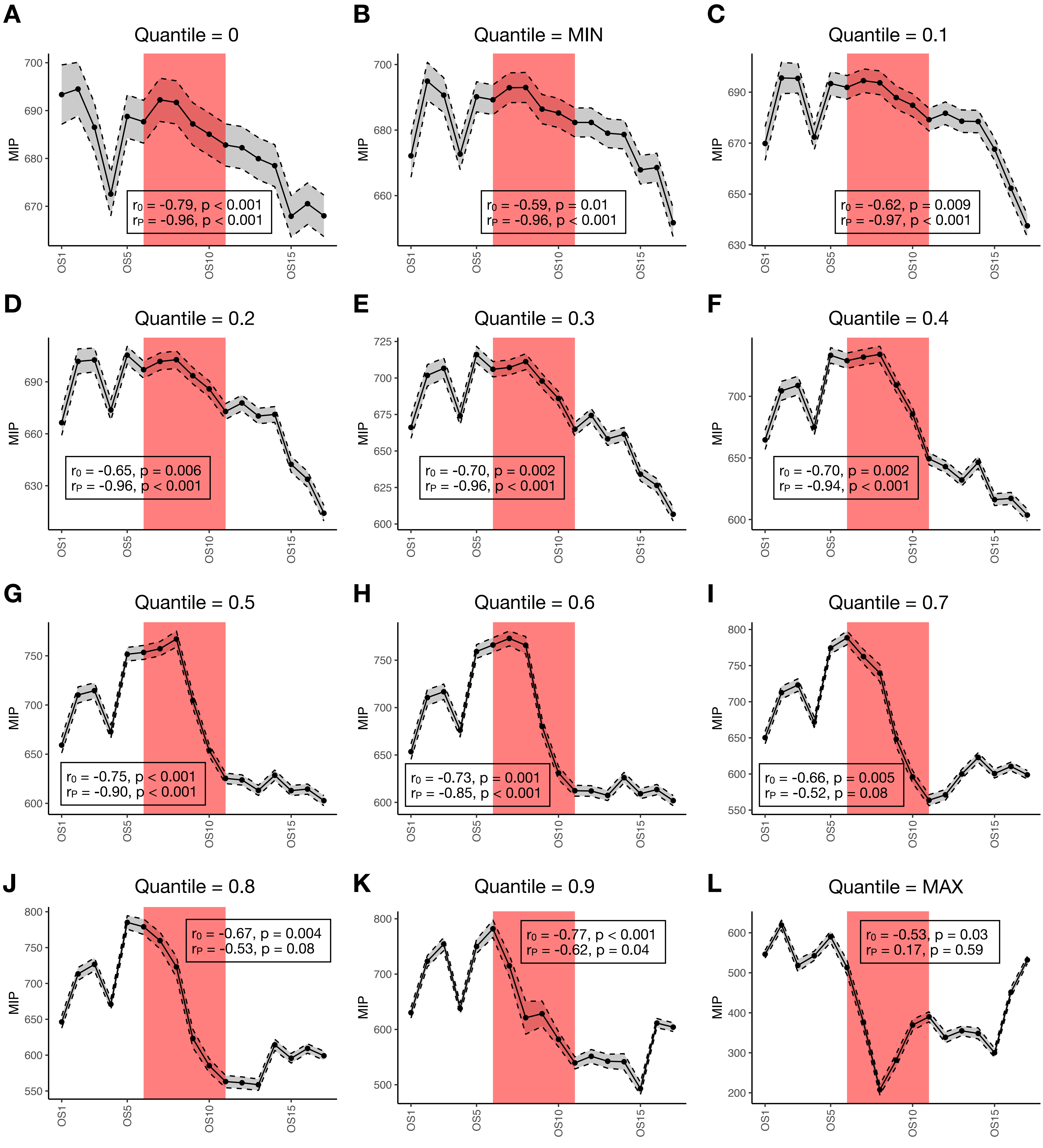

### supp_fig9.pdf

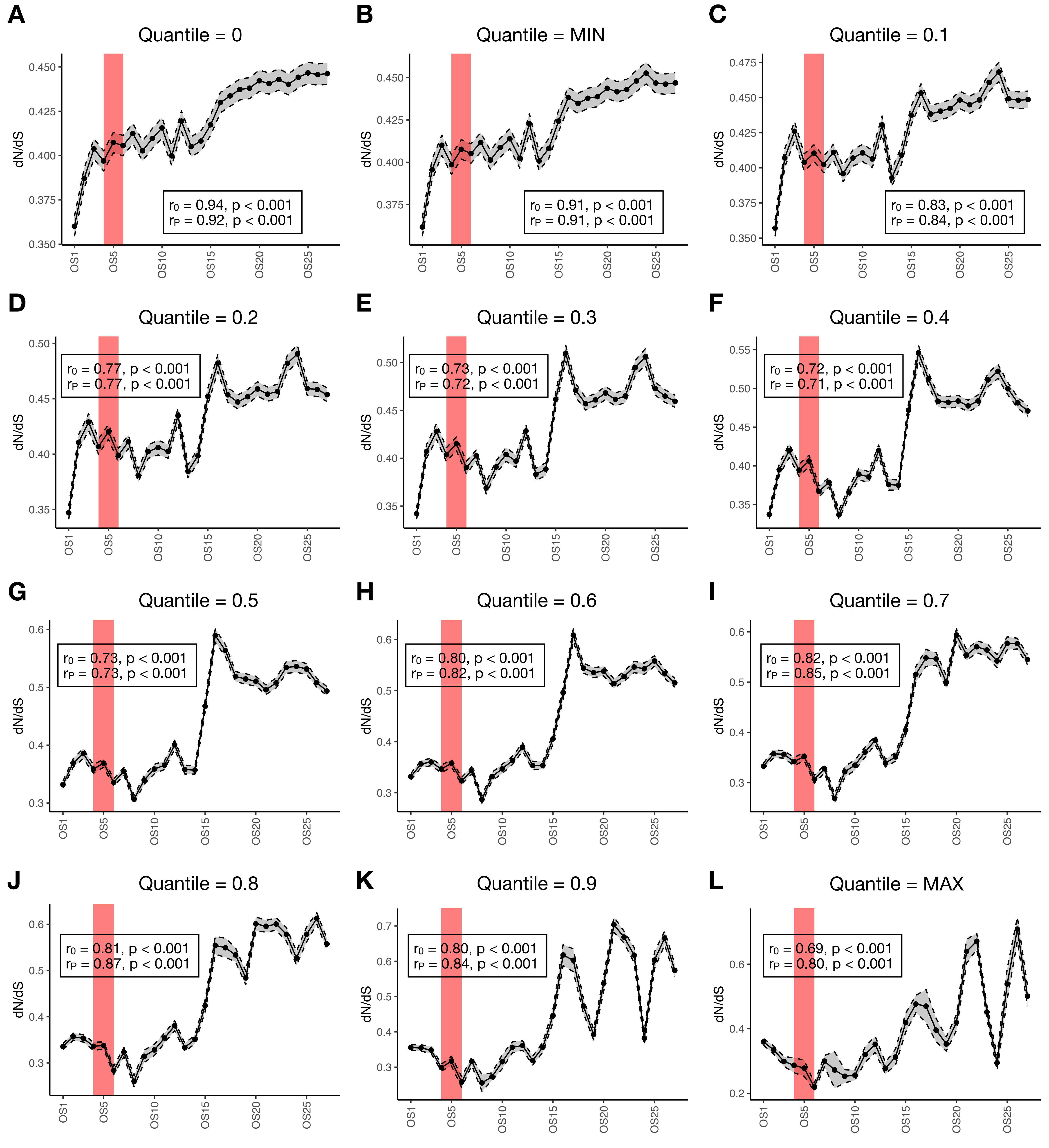

### supp_fig10.pdf

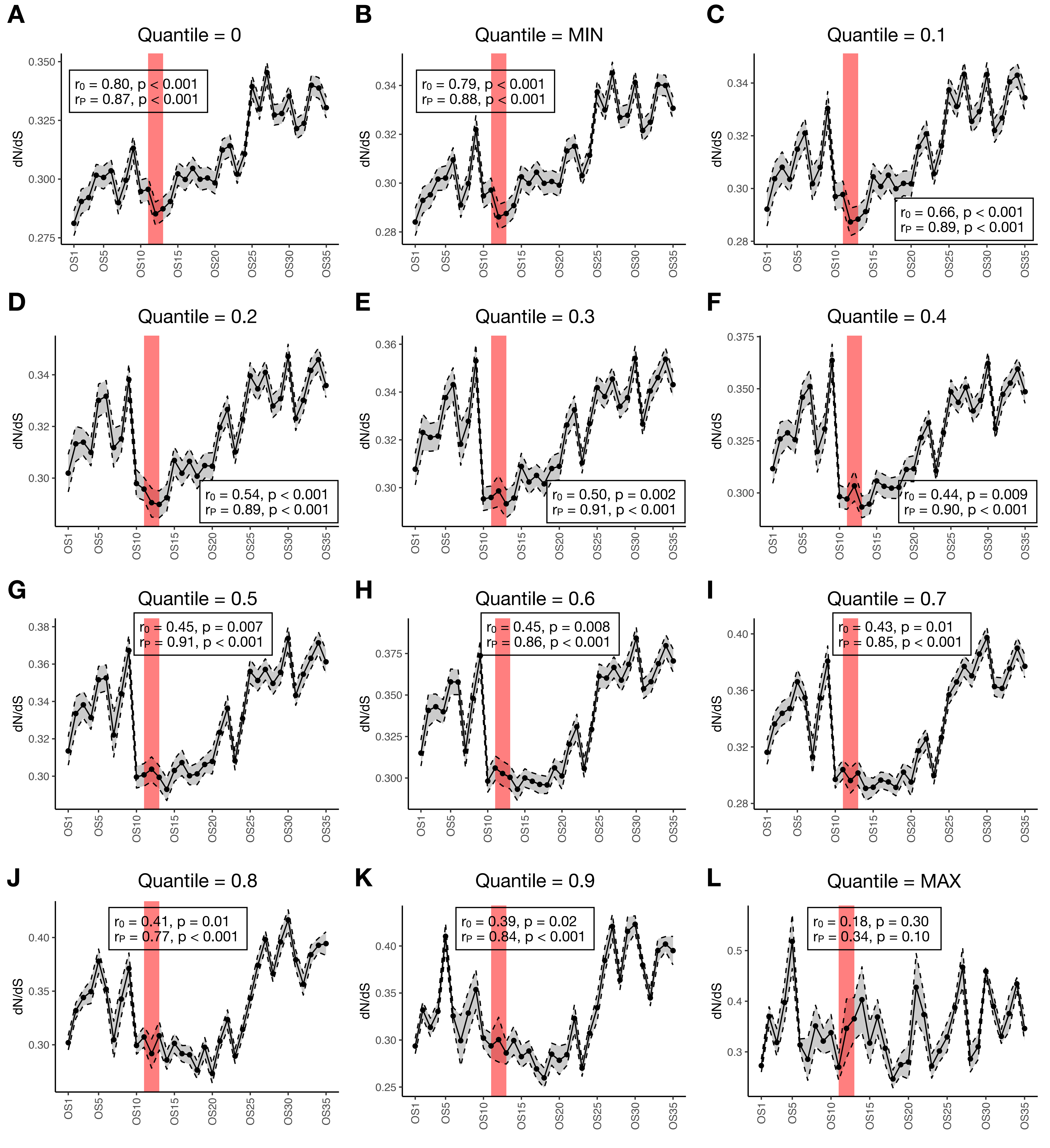

### supp_fig11.pdf

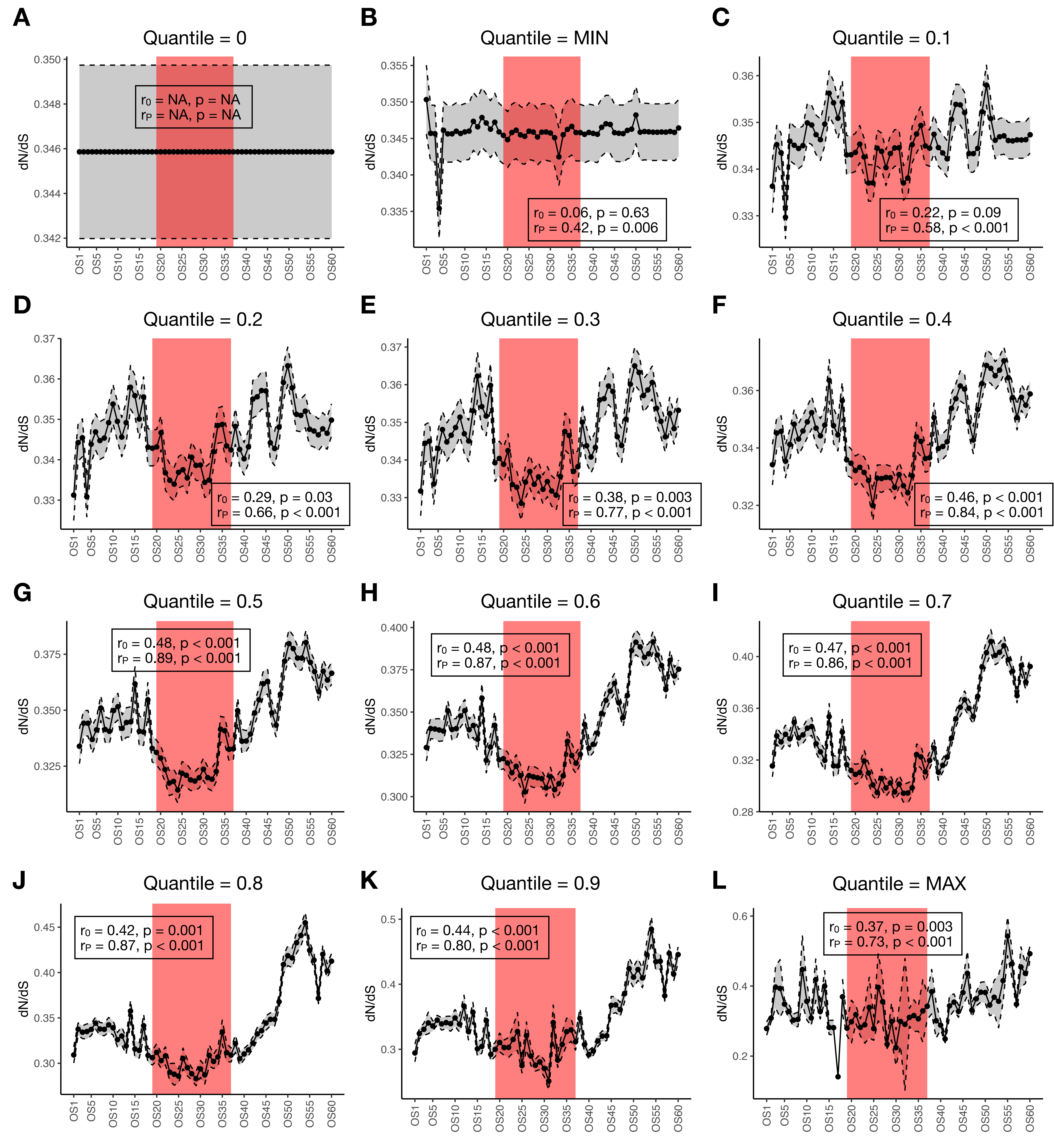

### supp_fig12.pdf

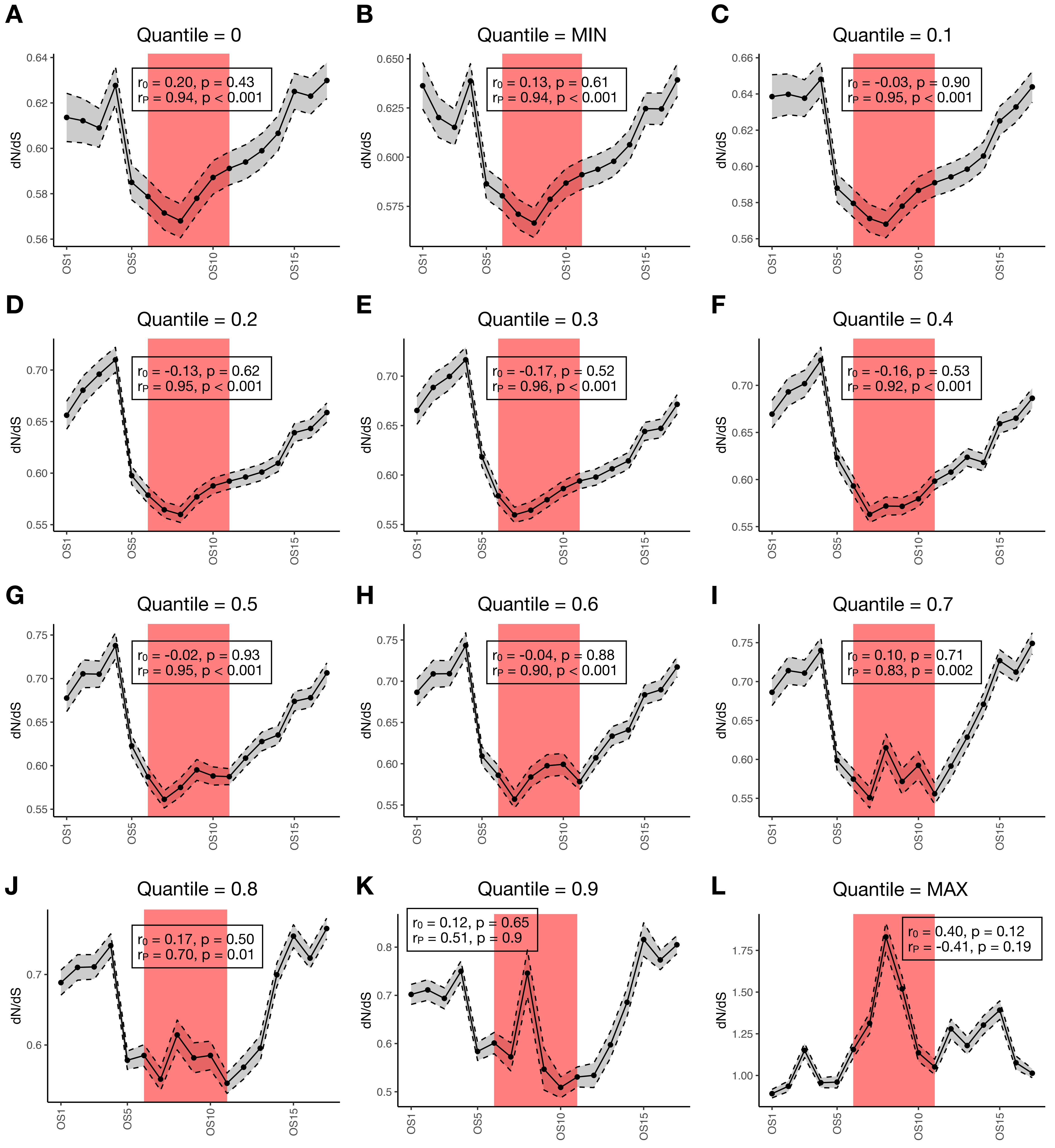

### supp_fig13.pdf

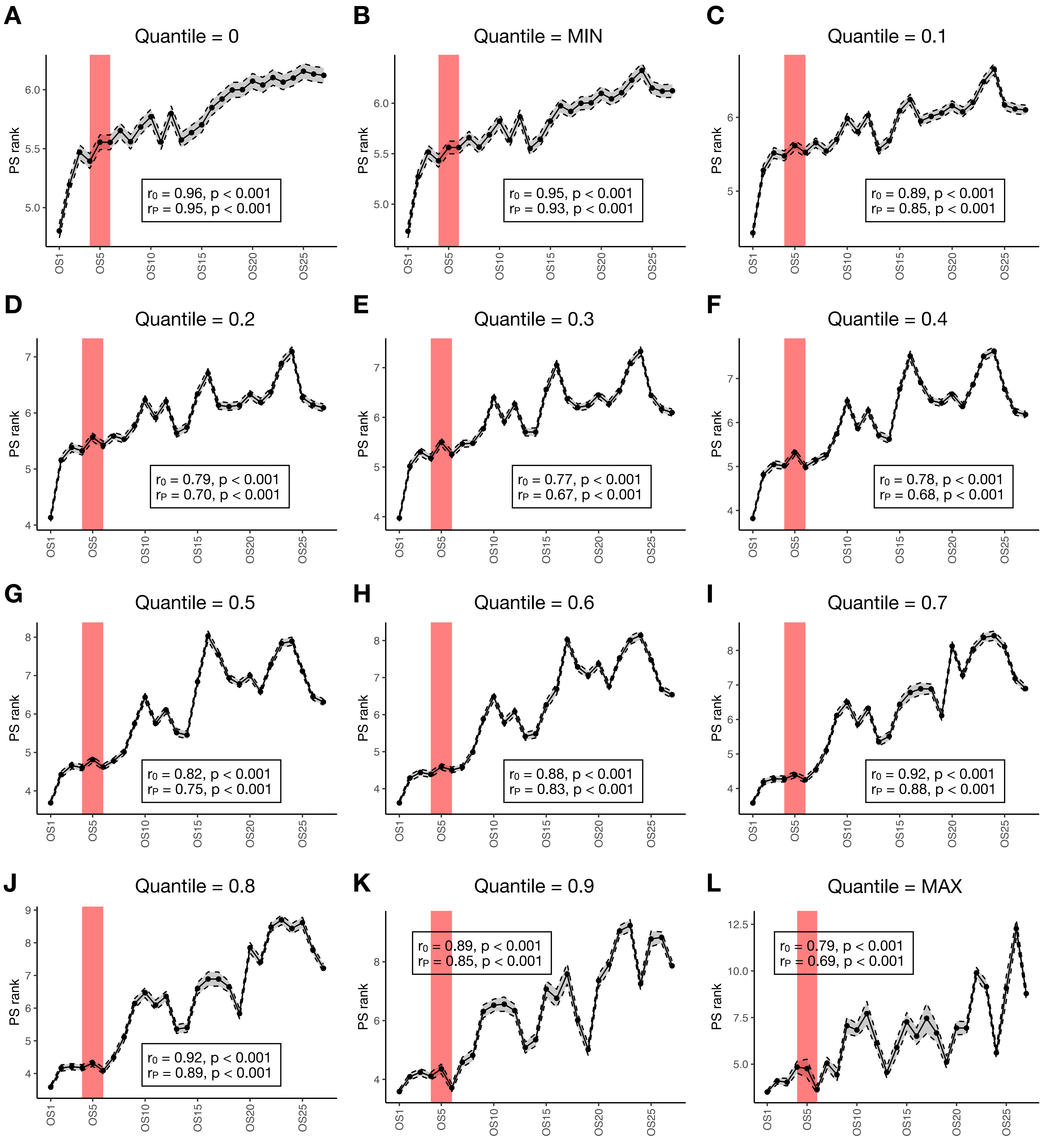

### supp_fig14.pdf

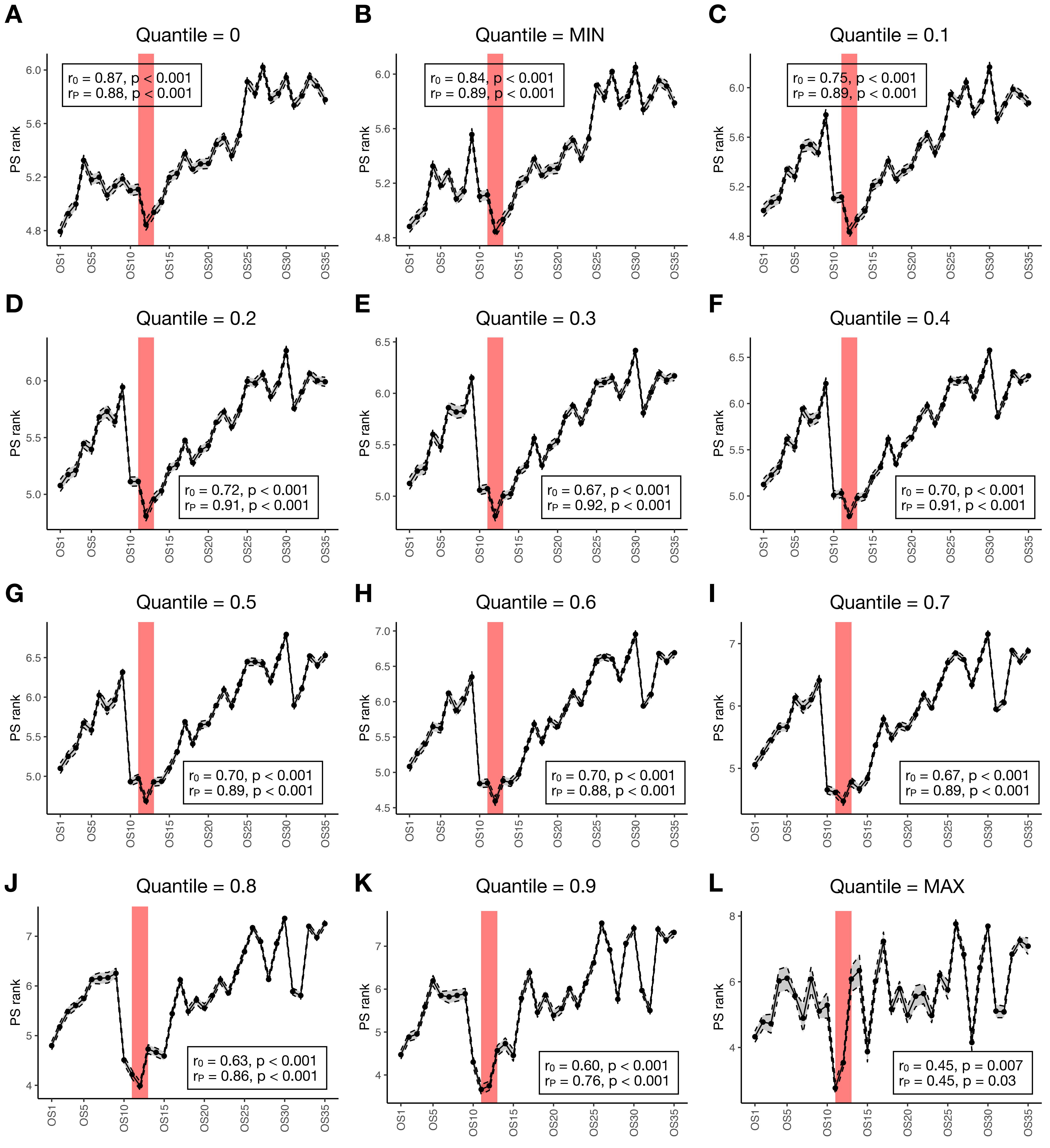

### supp_fig15.pdf

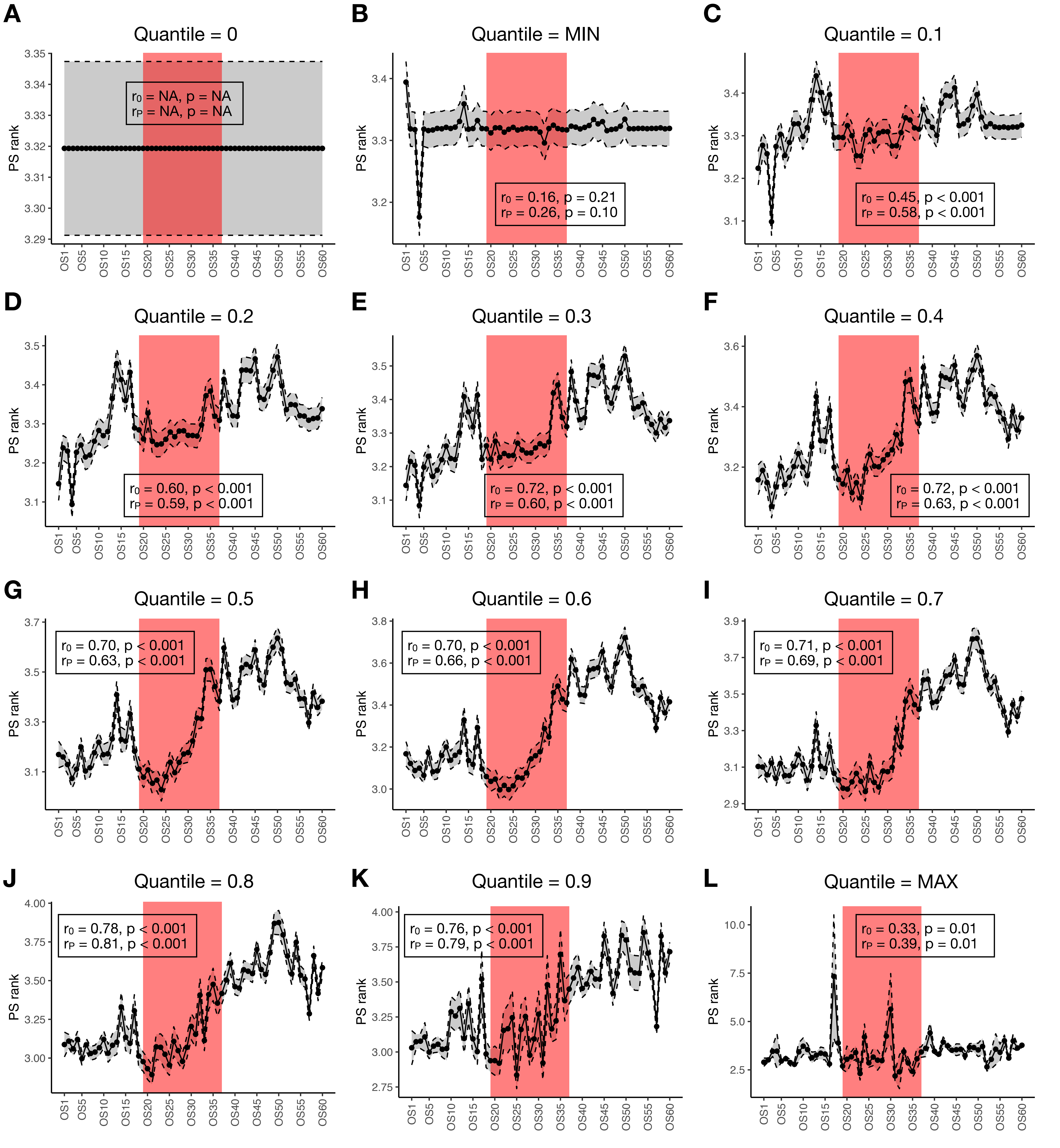

### supp_fig16.pdf

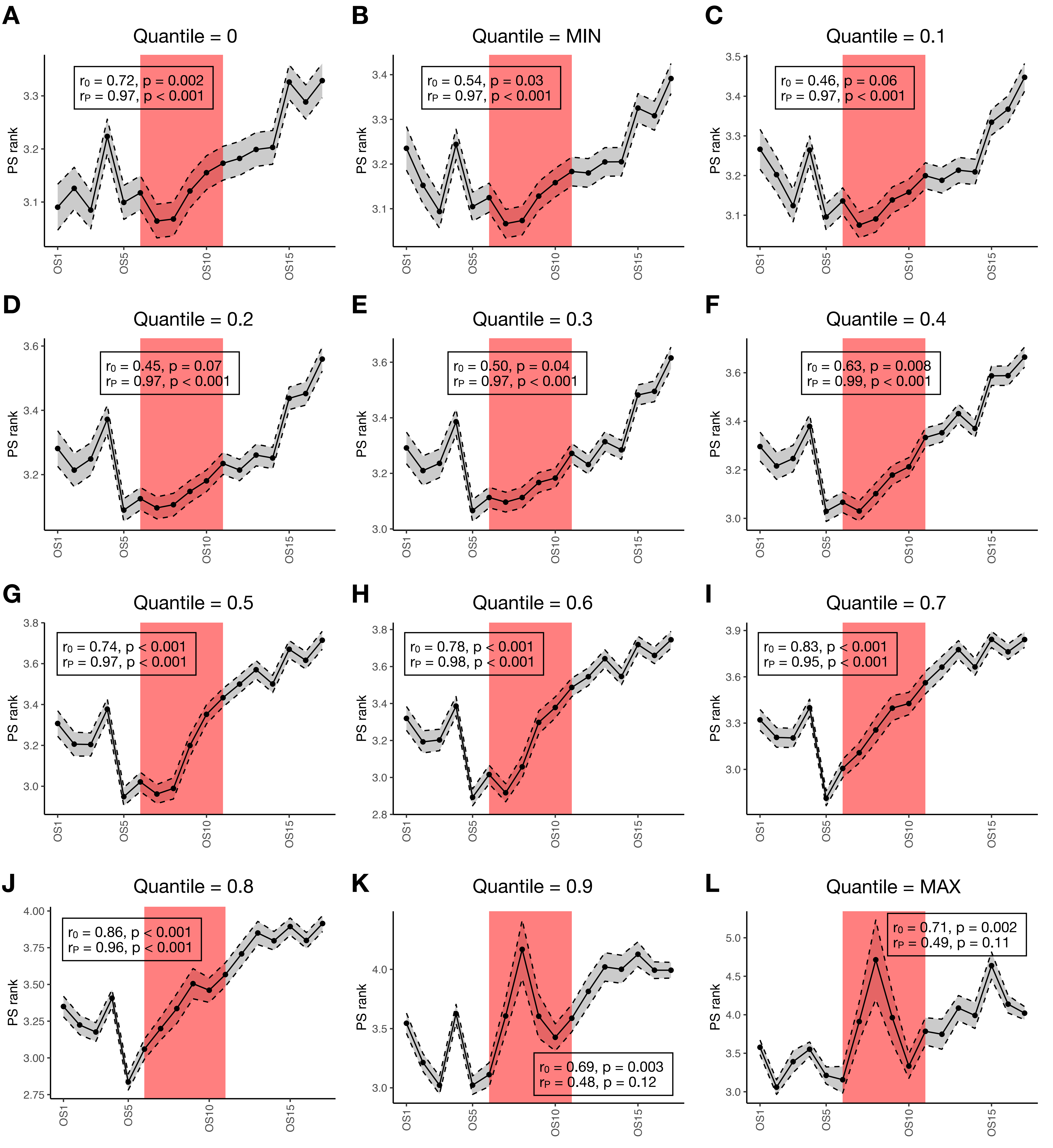
